# A Modeling Framework for Exploring Sampling and Observation Process Biases in Genome and Phenome-wide Association Studies using Electronic Health Records

**DOI:** 10.1101/499392

**Authors:** Lauren J. Beesley, Lars G. Fritsche, Bhramar Mukherjee

## Abstract

Large-scale association analyses based on observational health care databases such as electronic health records have been a topic of increasing interest in the scientific community. However, challenges of non-probability sampling and phenotype misclassification associated with the use of these data sources are often ignored in standard analyses. In general, the extent of the bias that may be introduced by ignoring these factors is not well-characterized. In this paper, we develop a statistical framework for characterizing the bias expected in association studies based on electronic health records when disease status misclassification and the sampling mechanism are ignored. Through a sensitivity analysis approach, this framework can be used to obtain plausible values for parameters of interest given results obtained from standard naïve analysis methods. We develop an online tool for performing this sensitivity analysis. Simulations demonstrate promising properties of the proposed approximations. We apply our approach to study bias in genetic association studies using electronic health record data from the Michigan Genomics Initiative, a longitudinal biorepository effort within Michigan Medicine.

## 1 Introduction

Genome-wide genotype data linked with electronic health records (EHR) are becoming increasingly available through biorepository efforts at academic medical centers, health care organizations, and population-based biobanks [1]. A common use of these linked data is to explore the association between a phenotype, *D*, with a risk factor of interest, *G*, after adjusting for potential confounders, *Z*. Analysis using a regression model for *D|G, Z* may be repeated for millions of risk factors or genetic variants with a given *D* of interest (as in a genome-wide association study (GWAS)) or for thousands of phenotypes derived from the content of the EHR with a given variant *G* of interest (as in phenome-wide association studies (PheWAS)). Association analyses embedded within large observational databases have gained popularity in recent years, and the use of and interest in such analyses continues to increase [1, 2, 3]. However, unlike curated and well-designed population-based studies, data obtained from large observational databases are often not originally intended for research purposes, and additional thought is needed to understand potential sources of bias. In this paper, we focus on the particular association setting with *D* being a single EHR-derived phenotype and *G* being a single genetic marker or a polygenic risk score, but the methods and conceptual framework developed in this paper can be applied quite broadly to general estimation problems using observational databases.

One potential source of bias in EHR-based studies is misclassification of derived disease status variables. Large-scale agnostic studies using EHR often define patient disease status (phenotypes) based on international classification of disease (ICD) codes or aggregates thereof called “PheWAS codes” or “phecodes” to define phenotypes in an automated and reproducible way [4]. In practice, EHR-derived phenotype definitions are used to represent the underlying ‘true’ disease status. However, these ICD-based phenotype classifications can be erroneous in capturing the ‘true’ disease status for a variety of reasons. For example, psychiatric diseases can be difficult to diagnose, and diagnosis can often be subjective [5]. ICD code-based diagnoses may be an incomplete representation of a patient’s health state, which may also be recorded in doctor’s notes and elsewhere in the EHR. In response to this problem, there exists an extensive literature on using other structured and unstructured content of the EHR to define phenotypes more accurately [6, 7, 8, 9, 10]. Additionally, human validation can be used to evaluate phenotyping algorithms [11]. These existing phenotyping approaches can be effective in reducing misclassification given the information available in the EHR, but even the most sophisticated phenotyping algorithms cannot capture diagnoses that were *never recorded* in any form in the EHR. Secondary conditions may not always be entered into the EHR, and symptoms occurring between visits may not always be reported. The EHR cannot adequately capture diseases that a patient had prior to entry into the EHR (outside the observation window). The chance of correctly capturing a disease is inherently dependent on the length of stay in the EHR or the observation/encounter process for a given patient. We often have a systematic source of mis-classification (that we will call “observation window bias”) due to a lack of comprehensiveness of the EHR in capturing diagnoses or medical care obtained from outside sources (e.g. at another health care center). Together, these various factors can lead to a potentially large degree of misclassification, particularly due to underreporting of disease.

Several authors have proposed statistical methods for addressing misclassification of binary phenotypes in EHR-based studies. The extent of misclassification can be described using quantities such as sensitivity and specificity, but these quantities can vary from population to population and from phenotype to phenotype [12]. Huang et al. [13] proposes a likelihood-based approach that integrates over unknown sensitivity and specificity values but requires some limited prior information about the sensitivity and specificity. Wang et al. [14] proposes an approach for incorporating both human-validated labels and error-prone phenotypes into the estimation, but this approach will not account for observation window bias. Duffy et al. [15] and Sinnott et al. [16] expand on results in the measurement error literature to relate parameters in the model for the true outcome with the model for the misclassified outcome, but Duffy et al. [15] focuses on outcome misclassification with *binary* risk factors, and Sinnott et al. [16] focuses on the setting in which the probability of having observed disease is explicitly modeled using a variety of information in the EHR. Additionally, all of these methods do not directly address the sampling mechanism.

In addition to potential bias due to misclassification of disease phenotypes, the mechanism by which subjects are selected into the dataset can sometimes result in biased inference when not handled appropriately. Complex sampling designs in an epidemiologic study can be addressed using survey design techniques if the sampling strategy is known. However, the probability mechanism for inclusion of a subject into a biorepository is not *a priori* fixed or defined. Interactions with the health care system are generated by the patient, and it can be difficult to understand the mechanism driving sampling as well as self-selection for donating biosamples, which may be related to a broad spectrum of patient factors including overall health. Several authors recommend adjusting for factors such as number of health care visits or referral status to better account for the sampling mechanism [17, 18]. Additionally, there is a belief in the literature that gene-related association study results may be less susceptible to bias resulting from patient selection [19]. This belief stems from the assumption that an individual genetic locus is not usually appreciably related to selection. However, bias due to genotype relationships with selection can still arise in certain settings [20]. Additionally, a popular topic in genetics-related research right now is the use of polygenic risk scores, which combines information from many genetic loci into a score to quantify a patient’s genetic risk for developing a particular disease [21, 22]. While it may be reasonable to assume that a specific genetic locus may have little association with selection, this assumption becomes more tenuous for an aggregate polygenic risk score with stronger association with the underlying disease and other factors related to selection.

As we will demonstrate, patient sampling can create substantial bias in estimating genetic associations using EHR data in the presence of disease status misclassification. Existing statistical methods for dealing with phenotype misclassification do not directly take into account the mechanism by which patients are sampled and *vice versa*. Additionally, standard association studies often do not account for either source of potential bias. It is important to understand the implication of these sampling and observation processes on results from naïve association analyses.

Formal characterization of the settings in which we can expect bias and their impact on resultant inference may help guide study design and analysis in the future. In this paper, we develop a statistical framework incorporating both disease misclassification and the patient selection mechanism. We use this framework to characterize the amount of bias expected in EHR-based association study results when misclassification and the sampling mechanism are ignored in the naïve and commonly-used analysis approach. We focus on the particular setting in which phenotypes are underreported (naturally occurring for EHR data due to limited observation window bias), but we also provide an extension allowing for bidirectional misclassification. The analytical expressions enhance our understanding of study design and phenotype characteristics this bias may depend on. Through a sensitivity analysis approach, this framework can also be used to obtain plausible values for parameters of interest given results from simpler analysis methods. Simulations demonstrate promising properties of the proposed approach in capturing the true bias. We provide an interactive online tool for performing these sensitivity analyses. We apply our approach to study bias in genetic association studies using EHR-derived phenotypes in the Michigan Genomics Initiative, a longitudinal biorepository effort within Michigan Medicine.

## 2 Model Structure

Let the binary variable *D* represent a person’s true disease status. Suppose we are interested in the relationship between *D* and an individual’s inherited genetic information, *G*, adjusting for additional person-level information, *Z*. We will call this relationship the *disease mechanism* as seen in Figure 1. In genetic association studies, *G* may represent a single SNP (single nucleotide polymorphism) or a polygenic risk score [21]. In principle, however, *G* can be any risk factor of interest. *Z* often contains information such as age, gender, and several principal components for the patient’s genome-wide data. In practice, we may have many diseases or genotypes of interest in an association study, but for now we will consider the simple setting with single specified *D* and *G*.

**Figure 1:**
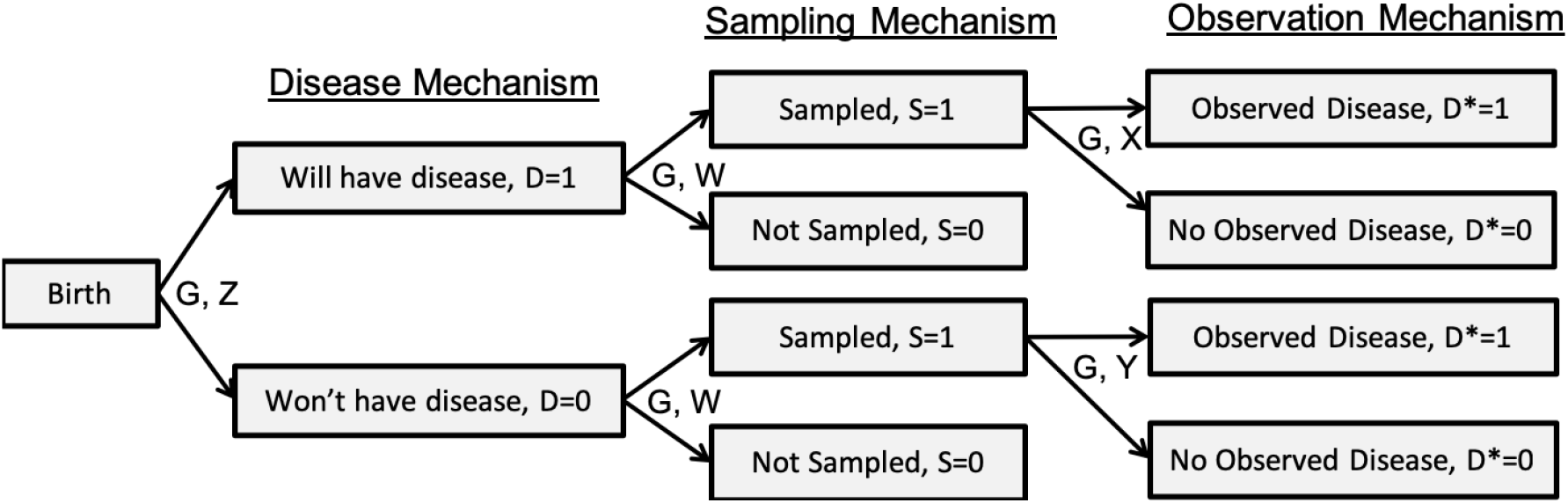
Diagram of the assumed data structure.

In order to make inference about *D*, we consider a large health care system-based database with the goal of making inference about the general population. Let *S* indicate whether a particular person in the population is selected into our dataset (for example, by going to a particular hospital and consenting to share a biosample for research), where the probability of a person in the population being included in the current dataset may depend on the underlying disease status, *D*, along with additional covariates, *W*, and perhaps even *G*. We may often expect the sampled and non-sampled subjects to have different rates of the disease, and other factors such as patient age, residence, access to care and general health state may also impact whether subjects are sampled into the study dataset or not. We will call this the *sampling mechanism*.

Instances of the disease are recorded in hospital or administrative records. We might expect factors such as the patient age, the length of follow-up, and the number of hospital visits to impact whether we actually *observe/record* the disease of interest for a given person. Define the *observed* disease status as *D\**. *D\** is a potentially misclassified version of *D*. We will call the mechanism generating *D\** the *observation mechanism*. There may be additional patient and provider-level predictors related to the true positive (sensitivity) and the false positive (1-specificity) rates, denoted *X* and *Y* respectively.

The diagram in Figure 1 shows the conceptual model structure, expressed as follows:

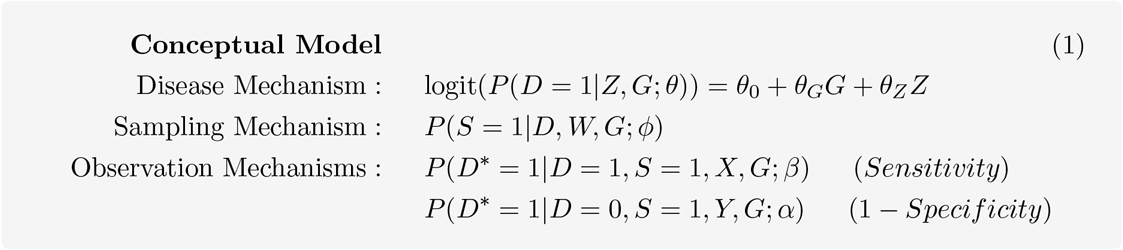

This framework allows for misclassification of the true disease status in either direction (subjects with the disease may be missed and subjects may be listed as diseased who aren’t). Moving forward, however, we will restrict our focus to the particular setting where disease status misclassification comes through underreporting. In other words, **we will assume perfect specificity** with *P* (*D\** = 1*|D* = 0*, S* = 1*, Y, G*; *β*) = 0 for all patients. This assumption may be reasonable for some EHR-derived phenotypes, particularly cancer, where we expect the rate of overdiagnosis of disease to be generally low. We consider the more general setting with imperfect specificity in detail in **Section S3**. We make the following assumptions:

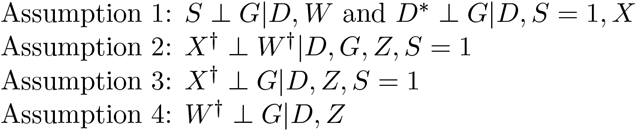

where *W*^†^ and *X*^†^ correspond to the predictors in *W* and *X not* included in *Z*.

The first assumption implies that ***G* is not an *independent* predictor of *S* and *D\****. This seems reasonable, especially when *G* represents a single SNP. A possible exception to this is in the case where *G* represents a rare, highly penetrant mutation. In this case, the SNP could conceivably be independently related to sampling. This is an important assumption, and our proposed methods may often under-estimate the bias of naïve analysis when this first assumption is violated. The second assumption states that the factors related to sampling and related to sensitivity that are not adjusted for in the disease model are independent given *D*, *G*, and *Z* on the sampled subjects. Importantly, this independence assumption conditions on *D*, *G*, and *Z*, and *Z* often includes common information such as patient age, gender, and the first few principal components of the genetic information. Conditionally, we may expect this assumption to be reasonable for many EHR settings. The third assumption implies that *G* is unassociated with the factors related to misclassification given *D* and *Z* on the sampled subjects. *X* is expected to contain information relating to a patient’s observation process, and this will generally not be driven by a patient’s genetic information given *Z* and *D*. These first three conditional independence assumptions, therefore, may often be reasonable in typical EHR data analysis settings.

The fourth assumption is the strongest, as it implies that ***G* is independent of factors related to sampling not included in the disease model given *D* and *Z***. Suppose, however, that sampling is related to a secondary disease, *D′*. This may often be the case for EHR data. If *D′* is independent of *G*, then this dependence will not create a problem and the fourth assumption will be satisfied. However, if *D′* is *independently related* to *G* (given *Z* and *D*), the fourth assumption is violated. An example of this would be the setting of pleiotropy, where a particular SNP may be related to multiple phenotypes, which could each contribute to sampling. This setting has been explored extensively in the literature on secondary analyses of case-control sampled data [e.g. 23, 24]. Importantly, the fourth assumption will be trivially satisfied if we include *D′* and other factors related to sampling as adjustment factors in the simple analysis model. Therefore, the proposed methods can be applied to characterize bias of naïve analysis even in the setting with sampling related to secondary diseases if we adjust for these secondary diseases in the simple analysis model. We discuss this challenge in more detail in **Section S5**.

## 3 Approximating Bias Resulting from Standard Analysis

### 3.1 Approximating the Bias

Suppose that *G, Z, X*, and *W* are observed if and only if *S* = 1. We further require that assumptions 1-4 hold. In standard analyses, we often fit a simple logistic regression for *D*|G, Z, S* = 1 (*analysis model*) with the goal of making inference about *θ* from the *target model* as follows:

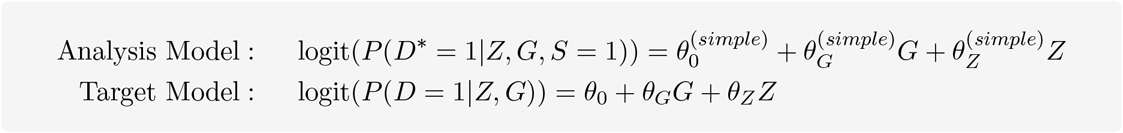

In general, *θ*_*G*_ and 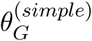 may be unequal. As shown in the **Supplementary Section S2**, we can relate the analysis model to the conceptual model in (1) as follows:

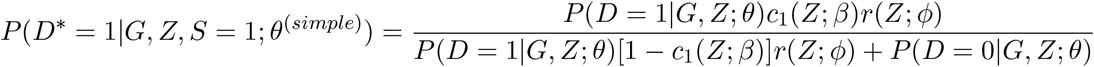

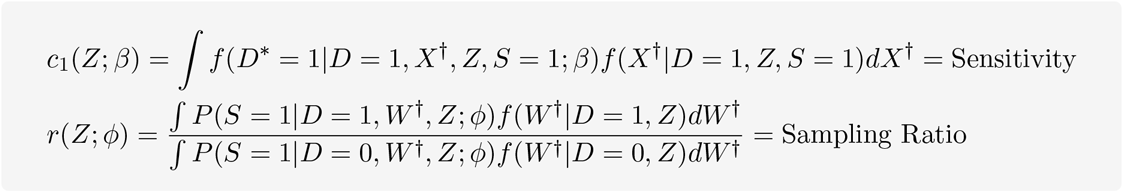

We can view *D\** as a noisy diagnostic for the true value of *D* with potentially imperfect sensitivity, where *c*_1_(*Z*; *β*) represents the average sensitivity of *D\** for *D* in the sampled subjects. We consider the more general setting with imperfect specificity in **Section** S3.1. *r*(*Z*; *φ*) represents the average sampling ratio comparing subjects with *D* = 1 and *D* = 0. These expressions may both depend on *Z*. However, *c*_1_(*Z*; *β*) and *r*(*Z*; *φ*) depend on distinct parameters, so they can vary independently conditional on *Z*. These functions depend on various covariate distributions, but we will *not* need to specify these distributions in practice.

We assume that covariates *Z* are centered so that they have mean zero and that the data are modeled as in (SuppEq. 1.1). Let 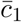 and 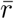 be the above expressions evaluated at 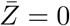. As shown in **Supplementary Section** S2.1, we use Taylor series approximation to obtain expressions for the intercept and coefficient for *G* in the *analysis model* as functions of parameters in the *target model*. We consider two different cases with two types of genetic variables *G*.

#### Case 1: *G* is a SNP

Suppose that *G* represents a single genetic locus or SNP (single nucleotide polymorphism), coded 0/1/2 to represent the number of copies of the minor allele for each patient at a bi-allelic locus. Let *MAF* denote the minor allele frequency in the *population*. We can express the *analysis model* parameters as a function of the *target model* parameters as follows:

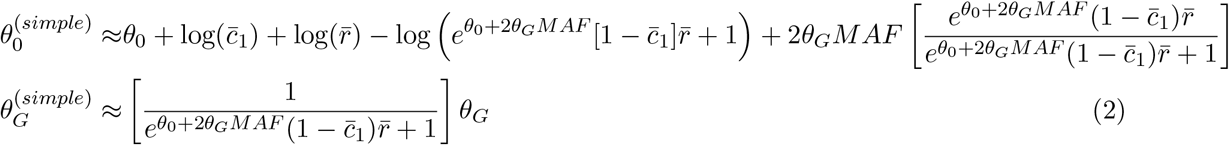

Here, we approximate the sample MAF with the population MAF as discussed in **Supplementary Section** S3.1.

#### Case 2: G is a Polygenic Risk Score

Suppose instead that *G* is a continuous predictor such as a polygenic risk score (as discussed in Dudbridge [25]) and suppose *G* has been centered to have mean zero. In this setting, we can express the *analysis model* parameters as follows:

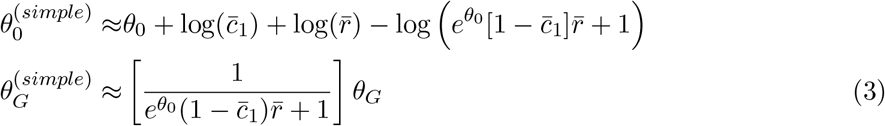

Simulations exploring the correspondence between the approximations in (2) and (3) and naïve estimated parameters can be found in **Figure S2**. These simulations indicate an excellent ability of these expressions to recover bias observed in simulated data.

### 3.2 Understanding the Structure of the Bias

We can use expressions (2) and (3) to develop some intuition for settings in which we expect greater or less relative bias when performing the “simple” routine analysis. Table 1 describes the general impact of the various model parameters on the bias of 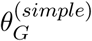, the parameter associated with *G* in the simple analysis model. As expected under assumptions 1-4, there is no bias in estimating 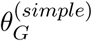 when we have perfect sensitivity. This may not be the case if one or more of these assumptions are violated. Suppose instead that 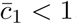. In this case, we expect bias in 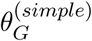, and the absolute bias will depend on the sensitivity, the disease sampling ratio 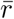, the population prevalence of the disease, the magnitude of *θ*_*G*_, and (in case 1) the *MAF* of the genetic locus of interest.

**Table 1:** Bias (relative and absolute) in target model parameters in the simple analysis

| Variable | Value | $G$ is a PRS*<br>Bias of $\theta_0^{(simple)}$ | $G$ is a PRS or SNP<br>Bias of $\theta_G^{(simple)}$ |
| --- | --- | --- | --- |
| Sensitivity ($\bar{c}_1$) | Decreases toward 0<br>No underreporting ($\bar{c}_1 = 1$)<br>Underreporting ($\bar{c}_1 < 1$) | Bias increases<br>Biased if $\bar{r} \neq 1$ or $r^{cc} \neq 1^\dagger$<br>Biased | Bias increases<br>No bias<br>Biased |
| Sampling ratio ($r$) | Increases toward ∞<br>Sampling independent of $D$ ($\bar{r} = 1$)<br>Sampling dependent on $D$ ($\bar{r} \neq 1$)<sup>†</sup> | Bias depends on $\theta_0, \bar{c}_1$<br>Biased if $\bar{c}_1 < 1$<br>Biased | Bias increases if $\bar{c}_1 < 1$<br>Biased if $\bar{c}_1 < 1$<br>Biased if $\bar{c}_1 < 1$ |
| Disease prevalence | Increases toward 1 | Bias depends on $\bar{r}, \bar{c}_1$ | Bias increases if $\bar{c}_1 < 1$ |
| Absolute effect of $G$ | Increases toward ∞ | Little or no added bias | Bias increases if $\bar{c}_1 < 1$ |

\*Direction of bias of $\theta_0^{(simple)}$ under case 1 ( $G$ is a SNP) depends on other parameters
<sup>†</sup> We consider sampling dependent on *observed* case-control status, $D^*$ , in **Section S4**.

Suppose we assume that the sampling ratio is 1 or 5, so diseased and non-diseased people are sampled at a 5:1 or 1:1 ratio on average respectively. Figure 2 shows the value for 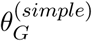 we may obtain if we fit a model for *D*|G, Z* using the sampled data with a minor allele frequency of 0.2 and true *θ*_*G*_ either 0.5 or 0.1. As the sensitivity decreases, the “simple” estimate of *θ*_*G*_ becomes increasingly biased (relative and absolute bias) toward the null. The level of bias depends strongly on the population prevalence of the disease. For less common diseases (e.g. less than 10% of the population), we may expect to observe relatively low relative bias in estimating *θ*_*G*_ even with low sensitivity for observing *D* in the sampled patients. Additionally, we expect the absolute bias to be generally small when we have lower true values for *θ*_*G*_. For moderate to large values of *θ*_*G*_, misclassification and sampling can have a substantial impact on the absolute bias for 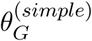. This provides some support to the notion that absolute bias in GWAS studies (where the *θ*_*G*_ values are expected to be generally small) may be less of a concern, particularly when we are studying a less common disease. However, if we are studying a disease that has higher population prevalence or larger values for *θ*_*G*_ (for example, a highly penetrant SNP), we may be more concerned about the potential for absolute bias induced by ignoring disease misclassification and/or sampling. As shown in **Table S1**, the bias has a more complicated relationship with disease prevalence and sensitivity in the presence of disease overreporting. We created an online RShiny tool called SAMBA-EHR (**S**ampling **A**nd **M**isclassification **B**ias **A**nalysis for EHR-based association studies) for exploring the impact of the various parameters on the bias for both the intercept and association parameters, available at http://shiny.sph.umich.edu/SAMBA-EHR/.

**Figure 2:**
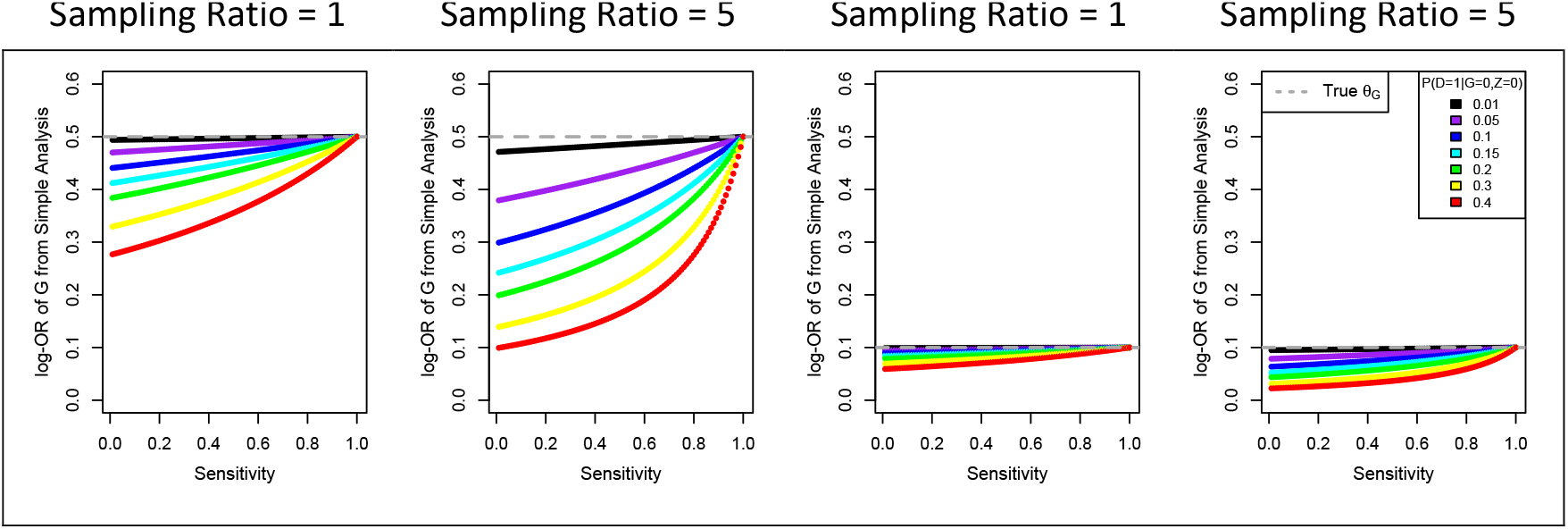
Bias of log-odds ratio for G when we model D*|G, Z, S = 1.

### 3.3 Sensitivity Analysis

An alternative use of the proposed model is in a sensitivity analysis in a reverse direction, where we can obtain reasonable values of *θ*_*G*_ across plausible values of 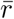 and 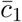 based on the parameter estimates from the “simple analysis.” Suppose we have an estimate for 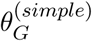 and that *G* is a single genetic locus. This estimate can be based on direct data analysis or can be obtained from published summary statistics. Using information from the population to obtain reasonable values for *θ*_0_, we can explore plausible values for *θ*_*G*_ by solving (2) for *θ*_*G*_. In practice, *θ*_0_ itself may not be known. In this setting, we propose performing the sensitivity analysis for a plausible window of *θ*_0_ using a rough estimate for the population prevalence, *P* (*D* = 1). Alternatively, suppose that 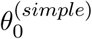 is also available. In this case, we can obtain *θ*_*G*_ by solving

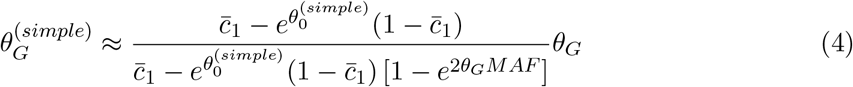

for *θ*_*G*_. In this case, predicted *θ*_*G*_ will be a function of 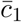 and *θ*^(*simple*)^ but not 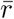. Similar expressions for predicting *θ*_*G*_ when *G* is a PRS can be found in **Section** S2.2, and the setting with imperfect specificity is considered in **Section** S3.2. In some settings, these expressions may have no solution for given values of 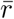 and 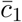. When this happens, it usually corresponds to a setting that is implausible given the observed data. For reasonable combinations of parameters, however, these expressions can be used in a sensitivity analysis after the primary analysis has been completed to explore the potential impact of ignoring misclassification and the sampling mechanism.

## 4 Simulations Evaluating Proposed Sensitivity Analysis

The usefulness of the expressions in (2) and (3) relies on the accuracy of the Taylor series approximations used to derive them. In **Section S6**, we demonstrate strong concordance between (2) and (3) and estimates from simulated data. One use of the approximations in (2) and (3) is to guide a sensitivity analysis after *θ*^(*simple*)^ has already been estimated, and we describe how this can be done in Section 3.3.

Here, we present results from a simulation study in which we compare the correspondence between true values of *θ*_*G*_ and values predicted by the sensitivity analysis in different settings. We will focus on the setting where *G* is a genotype on a single marker as this setting is more complicated than the PRS setting. A detailed description of the simulation procedure can be found in **Section S6**. Briefly, we simulate data under different combinations of 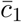, 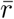, and *P* (*D* = 1*|G* = 0*, Z* = 0) (100 datasets each).

Figure 3 shows predictions for *θ*_*G*_ (from solving (4) for *θ*_*G*_) across several simulation settings. These predictions assume 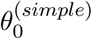 is available and do not depend on 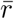. Each subplot corresponds to a different combination of true 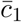 (denoted by the bolded boxplot) and *P* (*D* = 1*|G* = 0*, Z* = 0) for the setting where 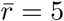. The boxplot for predicted *θ*_*G*_ corresponding to the true sensitivity (bolded) always includes the true value of *θ*_*G*_, and the corresponding median for the predicted *θ*_*G*_ is generally close to the true value of 0.5. These boxplots illustrate how a small population prevalence for the disease of interest is associated with greater robustness of inference (or good similarity between *θ*_*G*_ and 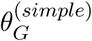 across sensitivity values. **Figures S3-S6** show predicted *θ*_*G*_ (from solving (2) for *θ*_*G*_) when we assume that a rough value for *θ*_0_ is available. These simulations again demonstrate the ability of the proposed sensitivity analysis to recover the true *θ*_*G*_ when evaluated at the true values for 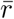 and 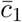. We emphasize, however, that we cannot use these sensitivity explorations to determine or estimate the “correct” values of 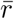 and 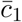 as they cannot be identified independently knowing only 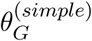 and *θ*_0_ or 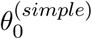. However, we can explore around reasonable values of 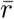 and 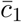 based on our understanding of the data to determine a range of plausible values for *θ*_*G*_.

**Figure 3:**
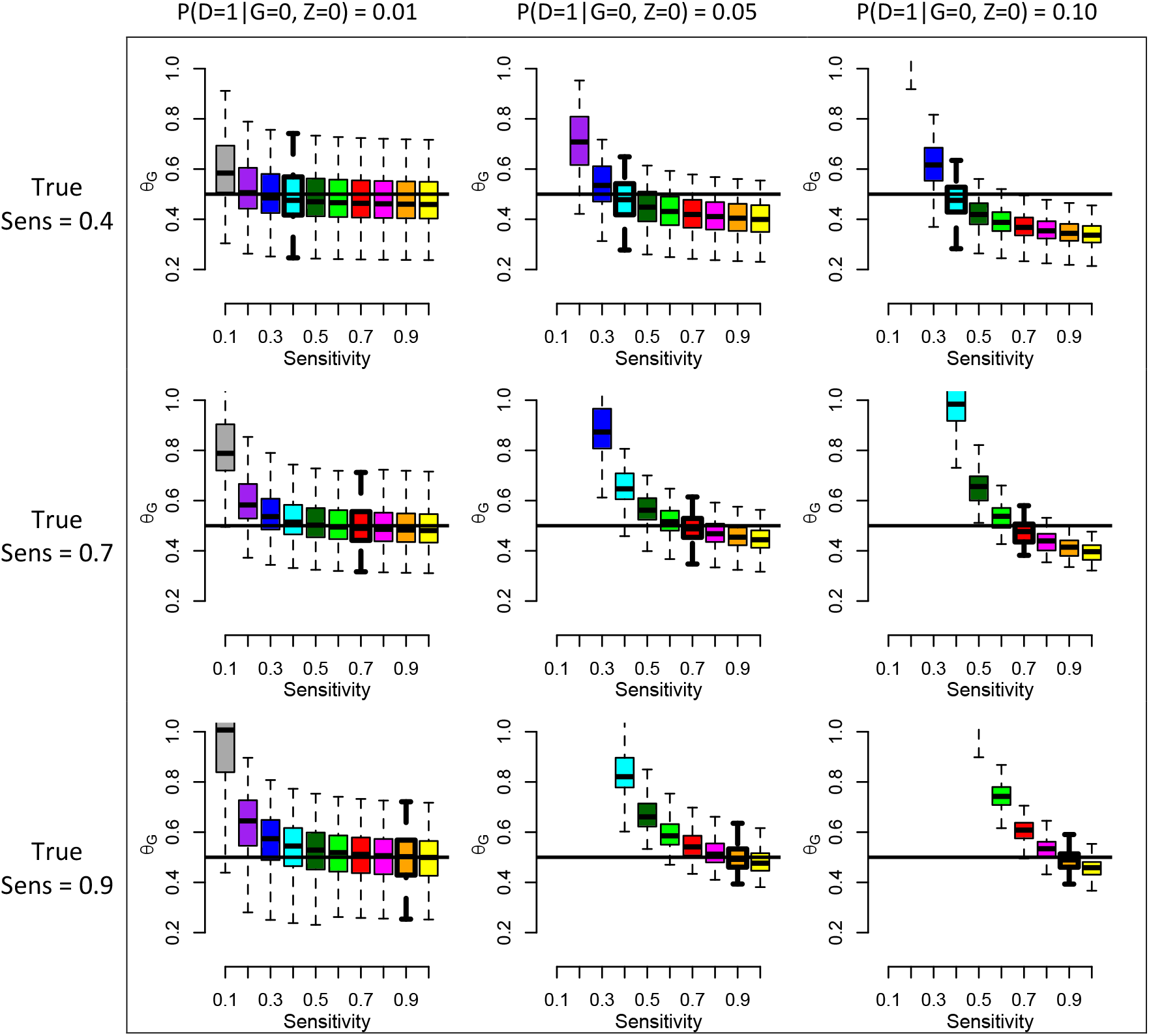
Predicted θ_G_ across different sensitivities using available 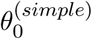. The black horizontal line indicates the true value of *θ*_*G*_ = 0.5. The bolded boxplot indicates the true sensitivity. Each boxplot represents predicted *θ*_*G*_ for 100 simulated datasets. True 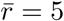

## 5 Bias Exploration in the Michigan Genomics Initiative

The Michigan Genomics Initiative (MGI) is a longitudinal biorepository effort within Michigan Medicine linked to EHR containing over 30,000 patients with matched genotype and phenotype information. We have strong reason to believe that sampling in MGI is somehow related to patient characteristics and, in particular, underlying disease status. Firstly, all patients in MGI are people who visited Michigan Medicine. Patients may go to the doctor when they feel sick or for other personal reasons, and these reasons are often unknown. Additionally, patients are largely recruited into MGI from the pool of subjects having surgery at Michigan Medicine. Patients in MGI tend to be sicker and have a greater number of diagnoses than the general Michigan population and even the Michigan Medicine population [1]. Due to its sampling mechanism, MGI is strongly enriched for cancer cases, as shown in Beesley et al. [1] and reproduced in **Table S2**. The large number of cancer cases makes MGI an attractive dataset for exploring associations between cancer diagnosis and genetics.

We are interested in associations between true underlying cancer status and genetic factors. When we have perfectly classified phenotypes (so there is no observation window bias and no under-or over-reporting of disease status), simple association analyses may produce reasonable results. However, when disease status is potentially misclassified, disease-related sampling 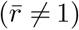 can result in biased estimates from the target model.

Previous analyses have explored associations between single genetic markers and polygenic risk scores with phenotypes for patients in MGI using simple analysis methods involving fitting a model for *D*|G, Z, S* = 1 [1, 21]. These analyses ignore the potential bias induced by the sampling and phenotype misclassification mechanisms. We are interested in evaluating the potential impact of the sampling and misclassification mechanisms on inference. Below, we perform the proposed sensitivity analyses to study to robustness of SNP-phenotype and PRS-phenotype associations in MGI for different assumed sampling and phenotype misclassification mechanisms. Since we are interested in cancer phenotypes, which may only rarely be incorrectly diagnosed, we assume perfect specificity for these analyses.

### 5.1 Genetic association analysis using individual markers

Following Beesley et al. [1], we perform a breast cancer GWAS using a cohort of over 38,000 patients in MGI. This analysis was performed on a matched subset of patients based on age and the first four principal components of the genome-wide data, and our results also adjusted for age and the principal components. The phenotype generation, locus pruning, and GWAS analysis were performed following Beesley et al. [1]. The breast cancer phenotype was defined using phecodes based on ICD codes using the R package PheWAS [4].

We first compare our simple GWAS results in MGI 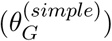 with reported associations from the NHGRI-EBI GWAS catalog (*θ*_*G*_), which combines results from meta-analysis of the largest and highest quality studies. We treat NHGRI-EBI GWAS catalog as a comparative gold standard. As shown in **Figure S7**, the association results using MGI data are generally similar to results in the GWAS catalog, but there are some specific SNPs for which the results differ. We focus our attention on 6 individual loci for which the GWAS catalog estimate (*θ*_*G*_) differs appreciably from the MGI estimate 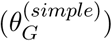. For this exploration, we fix *θ*_0_ = logit(0.124), using the US female population lifetime rate of breast cancer [1].

In Figure 4, we explore plausible values of *θ*_*G*_ in MGI across different potential sampling ratios and sensitivities. We obtain rough intervals for *θ*_*G*_ for fixed values of 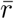 and 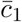 using the upper and lower 95% confidence interval limits of the MGI GWAS estimate for each locus. In **Supplementary Section S8**, we describe one approach for getting a sense of plausible values for the sampling ratio. As either the sensitivity or sampling ratio goes to 1, the interval shown in the figure gets closer to the 95% confidence interval for the corresponding GWAS estimate in MGI. We note that, in principle, we expect some bias in 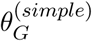 when we have imperfect sensitivity, but we see very little absolute bias when 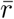 is small. This is primarily due to the small estimates for 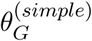. Bias in 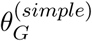 is expected to be proportional to the true value of *θ*_*G*_, and for small *θ*_*G*_, we may not see much absolute bias even for very low sensitivity when the sampling ratio is near 1. As the sampling ratio increases, however, the relative and absolute biases increas, and the predicted *θ*_*G*_ becomes more extreme. For the sake of comparison, we present an example for which the association in MGI and the GWAS catalog are nearly identical in the **Figure S8**. These results suggest that even GWAS results can potentially be impacted by sampling and misclassification. However, we note that comparison across multiple association tests within a GWAS study should have similar sampling ratios and misclassification rates. Therefore, we may have greater confidence in comparisons across association tests rather than in the absolute estimated effect for a particular association. We will explore this issue further in future work.

**Figure 4:**
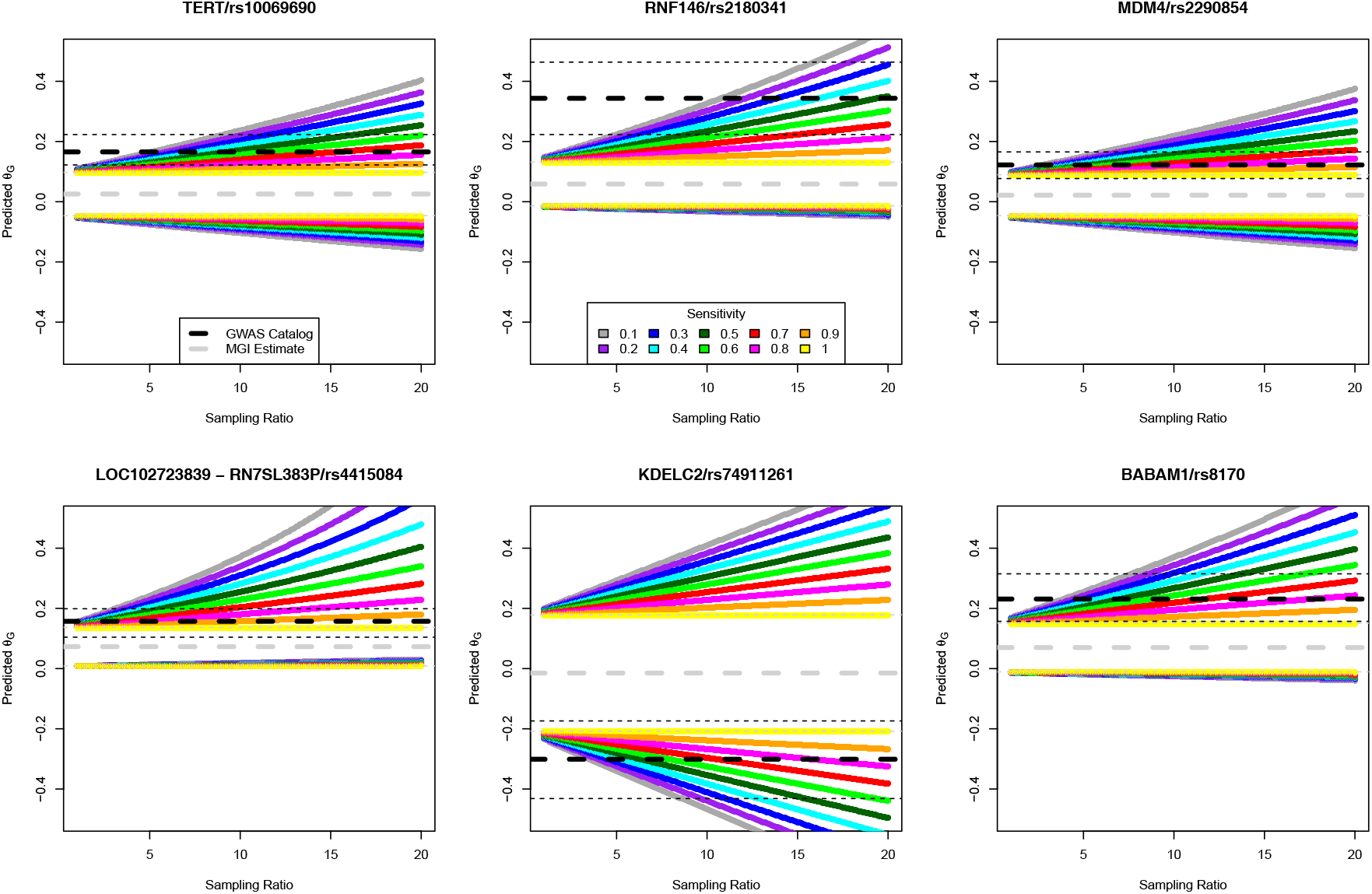
Plausible breast cancer log-odds ratios (θ_G_) in MGI for six SNPs known to be associated with breast cancer. The NHGRI-EBI GWAS catalog (Michigan Genomics Initiative) estimate and corresponding 95% confidence interval are shown in black (gray)

### 5.2 Association analysis with Polygenic Risk Scores

Previous work in Fritsche et al. [21] explored associations between polygenic risk scores (PRS) and their corresponding EHR-derived phenotypes in MGI for several cancer phenotypes. We are interested in exploring the robustness of these *published summary statistics* to different sampling and misclassification mechanisms for six different cancer diagnoses. We treat the published PRS association summary statistics (values listed in **Table S2**) as our 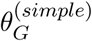 values, and we use the corresponding Michigan prevalence rates reported in Beesley et al. [1] to approximate *θ*_0_ for each of the six cancers of interest. We use the upper and lower confidence interval limits for 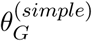 to create an interval for *θ*_*G*_.

Figure 5 shows the predicted interval for *θ*_*G*_ assuming different sampling ratios and sensitivities for each of the six selected cancers. Estimates for cancer of the bladder, non-Hodgkins lymphoma, and colorectal cancer are all very robust to different values for the sampling ratio and sensitivity. There are two primary reasons for this phenomenon. Firstly, the population prevalences of these three cancers are all low (less than 5%). As shown previously, we expect to see less relative bias in this setting. Secondly, the estimated association between the PRS and the disease is small in all of these cases. While the relative bias will not appreciably change as *θ*_*G*_ changes, the absolute bias will be small for small *θ*_*G*_. In contrast, breast and prostate cancer have high prevalences, and the corresponding PRS associations are strong. For these two PRS associations, our results suggest we may have appreciable relative bias if the sampling ratio is moderate to large and the disease outcome is at least moderately misclassified. In an extreme setting where the sensitivity is very low (e.g. 0.1) and the sampling fraction is fairly high (e.g. 10), we see predicted *θ*_*G*_ nearly doubles the observed 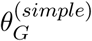 for both breast and prostate cancers. We explore plausible values for the sampling ratio for these cancers in MGI in **Figure S9**.

**Figure 5:**
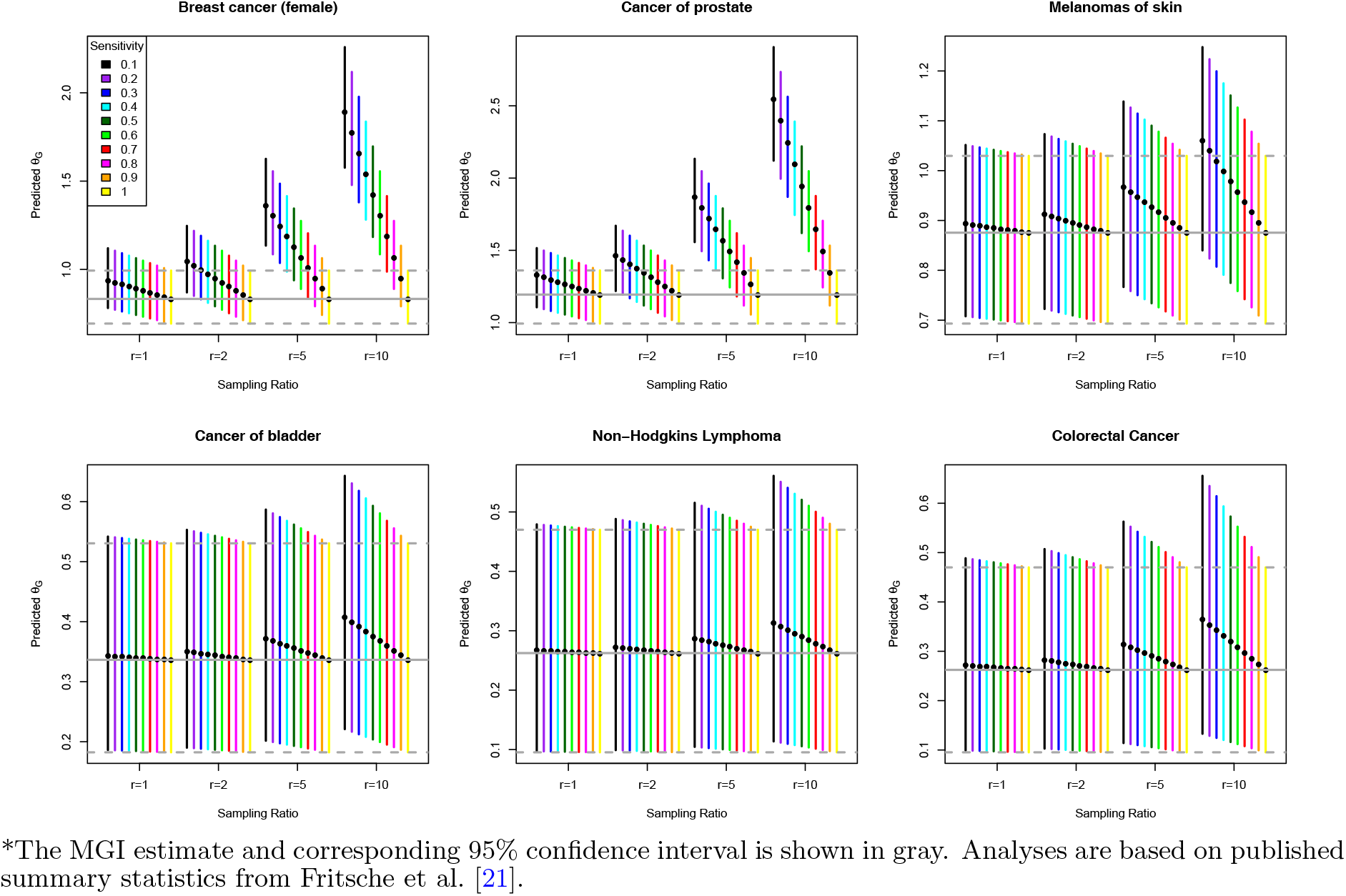
Sensitivity of PRS-phenotype associations for six different cancers*.

In all cases in Figure 5, the confidence interval for the PRS-phenotype association is far from zero, and adjustment for misclassification and sampling under our model would move estimated *θ*_*G*_ even farther from zero. Therefore, our general conclusions of a moderate or strong association between the PRS and the phenotype of interest remain robust across scenarios. However, we expect decreases in power for a test for 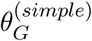 being nonzero when we have bias toward the null. We will evaluate properties of resulting tests and decision rules in future work exploring corrected estimation and inference techniques for *θ*_*G*_.

## 6 Discussion

The proposed modeling and sensitivity analysis framework allows us to explore the amount of bias we might expect in large association study results when we ignore issues of disease status misclassification and sampling related to disease and patient characteristics, as is often done in association studies using EHR-derived outcomes. Previous literature generally suggests that we may usually expect little bias in genotype-phenotype associations, and our statistical results lend credence to this belief when the disease of interest has low prevalence, say less than 10%, and sampling does not depend on the underlying disease status. When the disease of interest has higher prevalence in the population or the phenotype misclassification rates are higher, however, the sampling mechanism can have a substantial impact in biasing association results. Without any misclassification of the outcome, we do not expect much bias in log-odds ratio association parameters unless sampling is associated with both the underlying disease status and the covariate of interest given the adjustment factors. This is a property of the logistic regression modeling assumptions as has been explored in detail in the literature on secondary analysis of case-control sampled data and in **Section S5**, and modeling disease using different link functions may produce slightly different results [e.g. 23, 24].

We consider settings in which misclassification has perfect or imperfect specificity. Assuming perfect specificity (no overreporting of disease), we observe a higher potential for bias toward the null in the simple analysis for diseases with higher population disease prevalence. This assumption may be reasonable for many EHR-derived phenotypes, where the large part of misclassification is expected to be a result of underreporting of disease. In the setting of imperfect specificity, which may occasionally arise for diseases that are difficult to diagnose such as psychiatric disorders, we may have bias either toward or away from the null, and the general relationship between disease prevalence and bias is much less clear. The current exploration focuses on bias in effect estimates, but we may also be interested in p-values and hypothesis testing. We expect effect estimates biased toward the null to result in a loss of statistical power. We will explore the issues of Type I error and power in detail in a follow-up paper focused on estimation under the proposed model.

One advantage of the proposed modeling framework is that it does not require parametric modeling assumptions to be made for the sampling mechanism or the observation mechanisms (related to sensitivity and specificity). Instead, our results are guided by independence assumptions made on the relationships between drivers of these various mechanisms. A strong assumption made in the course of this paper is that the predictors related to sampling that are not included in the simple analysis model are independent of the genetic information of interest, conditional on the true disease status and adjustment factors *Z*. Since *Z* often contains age, gender, and several principal components of the genotype information, this may often be a reasonable assumption for EHR data. A challenging setting, however, is one in which sampling is related to a secondary disease *D′* that is *independently* related to *G* even adjusting for *D* and *Z* (perhaps due to pleiotropy). Our results can only be applied in this setting when simple analysis adjusts for any secondary diseases that are independently related to *G*. Secondary diseases independent of *G*, however, do not need to be adjusted for.

In this paper, we focus on estimation of a single genotype-phenotype association, but we are often interested in performing such estimation for many genotype-phenotype associations. A natural question is the extent at which this bias impacts comparison across estimated parameters in a large association study. In the case of GWAS, we perform association tests across many genotypes for a single phenotype. Here, the sampling ratio, sensitivity, and specificity are primarily properties of the particular disease we are interested in, and we do not expect these values to change much across the various association tests. In contrast, association tests in a PheWAS consider many different disease phenotypes. In this setting, we expect the sensitivity, specificity, and sampling ratios to differ across phenotypes, and accounting for differential bias toward the null across the various association tests may be of particular importance. There may be an opportunity to incorporate additional information such as the genetic architecture and disease heritability into the assessment of comparative bias.

The proposed framework can be useful for guiding analyses exploring sensitivity to violations of the common implicit assumptions of no outcome misclassification and ignorable sampling in EHR-based association studies. In future work, we will develop statistical methodology to perform parameter *estimation* under the proposed conceptual model. As part of the current work, we have developed an online tool called SAMBA-EHR (**S**ampling **A**nd **M**isclassification **B**ias **A**nalysis for EHR-based association studies) available at http://shiny.sph.umich.edu/SAMBA-EHR/. This will allow the proposed methods to be easily implemented in practice as a part of routine sensitivity explorations for association studies using EHR-derived outcomes.

## 7 Acknowledgments

The authors would like to thank Chad Brummet, Goncalo Abecasis, and Sachin Kheterpal along with the large group of collaborators at Michigan Genomics Initiative and also the MGI participants for generously donating their biosamples for research. This work was supported by NSF grant DMS1712933, The University of Michigan Comprehensive Cancer Center core grant supplement 5P30-CA-046592, and The University of Michigan precision health award U063790. The authors acknowledge the University of Michigan Medical School Central Biorepository for providing biospecimen storage, management, and distribution services in support of the research reported in this publication. *Dr. Lauren J. Beesley* wrote the manuscript and developed the methodology and the online SAMBA-EHR application. *Dr. Lars G. Fritsche* helped edit the paper and facilitated data acquisition and interpretation. *Dr. Bhramar Mukherjee* wrote the manuscript and provided key guidance on the methodology and manuscript.

