## Supplementary Materials for "A Modeling Framework for Exploring Sampling and Observation Process Biases in Genome and Phenome-wide Association Studies using Electronic Health Records"

May 14, 2019

### S1 Modeling and Assumptions

In the main paper, we propose the following conceptual model:

$$
\begin{aligned}
&\textbf{Conceptual Model} && && \text{(SuppEq. 1.1)}\\
&\text{Disease Mechanism :} && \text{logit}(P(D=1|Z,G;\theta)) = \theta_0 + \theta_G G + \theta_Z Z &&\\
&\text{Sampling Mechanism :} && f(S|D,W,G;\phi) &&\\
&\text{Observation Mechanisms :} && f(D^*|D=1,S=1,X,G;\beta) && \text{[Sensitivity Model]}\\
& && f(D^*|D=0,S=1,Y,G;\alpha) && \text{[1-Specificity Model]}
\end{aligned}
$$

This model allows for complicated sampling and observation mechanisms. Suppose our scientific interest is in the model for $D|G,Z$ and, in particular, the coefficient related to $G$. In practice, however, we do not fit the model in SuppEq. 1.1. We consider two general analysis models. In the first, we suppose we fit a model for $D^*|Z,G,S=1$ ignoring the potential misclassification and selection mechanisms. In the second analysis model, we suppose we fit a model for $D^*|Z,G,W,S=1$ ignoring the misclassification but adjusting for factors related to the sampling mechanism. Here, we clarify the various analysis and target models:

$$
\begin{aligned}
\text{Analysis Model :} \quad & \text{logit}(P(D^*=1|Z,G,S=1)) = \theta_0^{(simple)} + \theta_G^{(simple)} G + \theta_Z^{(simple)} Z\\
\text{Target Model :} \quad & \text{logit}(P(D=1|Z,G)) = \theta_0 + \theta_G G + \theta_Z Z \qquad \text{(SuppEq. 1.2)}
\end{aligned}
$$

Ideally, $\theta_G^{(simple)}$ and $\theta_G$ would be the same, but in practice this may not always be the case. Our strategy is to relate the parameters in the *analysis* models to the parameters in the *target* model. Here, we suppress the dependence of these distributions on parameters $\theta, \phi, \beta, \alpha$ and $\theta^{(simple)}$ in the notation. First, we consider different types of assumptions we can make on the relationships between $W$, $X$, $Y$, and $G$.

#### Potential independence assumptions

Define $W^\dagger$, $X^\dagger$, and $Y^\dagger$ to be the predictors in W, X, and Y not included in Z. Below, we list four different assumptions we make on these quantities.

Assumption 1: $S \perp G|D,W$ and $D^* \perp G|D,S=1,X,Y$ The former assumption states that G is not independently related to sampling given $D$ and $W$, so $f(S|D,W,G;\phi) = f(S|D,W;\phi)$. The latter assumption states that $G$ is not independently related to $D^*$ given $D$, $X$, and $Y$ in the sampled subjects, so $f(D^*|D=1,S=1,X,G;\beta) = f(D^*|D=1,S=1,X;\beta)$ and $f(D^*|D=0,S=1,Y,G;\alpha) = f(D^*|D=0,S=1,Y;\alpha)$

Assumption 2: $X^\dagger \perp Y^\dagger \perp W^\dagger|D,G,Z,S=1$ This assumption states that different, independent factors are related to the sensitivity and specificity mechanisms for observing $D$ conditional on $D$, $G$, $Z$, and $S=1$. We further assume that these are independent of $W$ on the sampled subjects given $D$, $G$, and $Z$. In EHR data, we might expect factors such as the length of follow-up, patient age, and the number of visits to be important factors in the sensitivity model. It seems reasonable that specificity (or the true negative probability) will be related to different factors. Factors such as age might often be related to both selection and disease misclassification, but age would often be included in $Z$. Critically, this independence assumption conditions on $D$, $G$, and $Z$, making this assumption much more plausible. For example, sampling may depend on another disease related to $D$, $Z$, and $G$. However, conditional on these variables, it seems plausible that $W$ is independent of $X$ and $Y$.

Assumption 3: $X^\dagger, Y^\dagger \perp G|D, Z, S = 1$ This assumption states that $X$ and $Y$ are independent of $G$ given $D$ and $Z$. This assumption seems reasonable as we do not expect factors related sensitivity/specificity to be associated with $G$ given $Z$, which may contain age, gender, and the principal components of the genetic information. This assumption seems particularly reasonable when $G$ represents a single SNP. However, there may be some settings where this is a **strong assumption**.

(Strong) Assumption 4: $W^\dagger \perp G|D, Z$ This assumption states that $W$ is independent of $G$ given $D$ and $Z$. This can be a **strong assumption** in some settings. Suppose, for example, that $G$ is a SNP that is independently related to two diseases, $D$ and $D'$, and that sampling is related to $D'$. In this setting, we will not have conditional independence between $G$ and $W$ given $D$ and $Z$. However, if $G$ is a SNP, the dependence between $G$ and $W$ may be so weak as to make the independence assumption reasonable in practice. If $G$ is a PRS independently related to $D'$, however, the *conditional* independence assumption bears additional thought.

Note on assumptions: Suppose that $W^\dagger$ is empty. In other words, assume all predictors in $W$ are included in $Z$. In this case, **Assumption 4** is trivially satisfied. In this cases, the analysis model would be a logistic regression including $W$ linearly. As shown in **Section S5**, this strategy is expected to have little bias in estimating $\theta_G$ when $W$ is independent of $G$ given $Z$, and there may be some settings where this strategy produces little bias even when $W$ is associated with $G$ given $Z$.

### S2 Bias and sensitivity analysis under perfect specificity

#### S2.1 Bias expressions

We will first assume that we fit analysis model 1 as in SuppEq. 1.2. However, we restrict our focus to the setting where we have perfect specificity, so $P(D^* = 1|D = 0, Y, S = 1; \alpha) = 0$ for all patients. This may be the most common setting when we are considering EHR data. We can write the analysis model as follows:

$$f(D^*|G, Z, S = 1) \propto \sum_d \int f(D^*|D = d, X, Y, G, S = 1) f(X^\dagger, Y^\dagger|D = d, G, Z, S = 1, W^\dagger)$$
$$\times P(S = 1|D = d, W, G) f(W^\dagger|D = d, G, Z) P(D = d|G, Z) dX^\dagger dY^\dagger dW^\dagger$$

Under **Assumptions 1-4**, we have that

$$f(D^*|G, Z, S = 1) \propto$$
$$P(D = 1|G, Z) \left[\int f(D^*|D = 1, X, S = 1) f(X^\dagger|D = 1, Z, S = 1) dX^\dagger\right] \left[\int P(S = 1|D = 1, W) f(W^\dagger|D = 1, Z) dW^\dagger\right]$$
$$+ P(D = 0|G, Z) \left[\int f(D^*|D = 0, Y, S = 1) f(Y^\dagger|D = 0, Z, S = 1) dY^\dagger\right] \left[\int P(S = 1|D = 0, W) f(W^\dagger|D = 0, Z) dW^\dagger\right]$$

Define

$$c_1(Z; \beta) = \int f(D^* = 1|D = 1, X^\dagger, Z, S = 1) f(X^\dagger|D = 1, Z, S = 1) dX^\dagger$$
$$r(Z; \phi) = \frac{\int P(S = 1|D = 1, W^\dagger, Z) f(W^\dagger|D = 1, Z) dW^\dagger}{\int P(S = 1|D = 0, W^\dagger, Z) f(W^\dagger|D = 0, Z) dW^\dagger}$$

Suppose we view $D^*$ as a noisy test for the true value of $D$ with potentially imperfect sensitivity and perfect specificity. We can view $c_1(Z)$ as the sensitivity of $D^*$ for $D$ in the sampled subjects averaged across the distribution of $X^\dagger$. We can view $r$ as the sampling ratio with respect to $D$, averaged across the distribution of $W^\dagger$. We note that $c_1(Z)$, and $r(Z)$ are functions of distinct parameters, so they can vary independently conditional on the observed data. Using this notation, we can rewrite the above expressions as

$$P(D^* = 1|G, Z, S = 1) \propto \left[\int P(S = 1|D = 0, W^\dagger, Z) f(W^\dagger|D = 0, Z) dW^\dagger\right] [P(D = 1|G, Z) c_1(Z) r(Z)]$$
$$P(D^* = 0|G, Z, S = 1) \propto \left[\int P(S = 1|D = 0, W^\dagger, Z) f(W^\dagger|D = 0, Z) dW^\dagger\right]$$
$$\times [P(D = 1|G, Z)(1 - c_1(Z)) r(Z) + P(D = 0|G, Z)]$$

Suppose we use logistic regression to model $D^*|G, Z, S = 1$ as in (SuppEq. 1.1) analysis model 1. We have

$$\begin{aligned}
&\text{logit}\left(P(D^* = 1|G, Z, S = 1)\right) \\
&= \log\left(\frac{P(D = 1|G, Z) c_1(Z) r(Z)}{P(D = 1|G, Z)(1 - c_1(Z)) r(Z) + P(D = 0|G, Z)}\right) \\
&= \log\left(\frac{e^{\theta_0 + \theta_G G + \theta_Z Z} c_1(Z) r(Z)}{e^{\theta_0 + \theta_G G + \theta_Z Z}(1 - c_1(Z)) r(Z)}\right) \\
&= \theta_0 + \theta_G G + \theta_Z Z + \log\left(c_1(Z)\right) + \log\left(r(Z)\right) - \log\left(e^{\theta_0 + \theta_G G + \theta_Z Z}(1 - c_1(Z)) r(Z) + 1\right)
\end{aligned}$$

We now approximate the above expression using a first order Taylor Series approximation with respect to $Z$ and $G$, where $\bar{Z}$ represents the mean of $Z$ and $\bar{G}$ represents the mean of $G$ in the sample. Let $\bar{c}_1 = c_1(\bar{Z})$, and $\bar{r} = r(\bar{Z})$. We can write

$$\text{logit}\left(P(D^* = 1|G, Z, S = 1)\right) \approx \theta_0 + \theta_G G + \theta_Z Z + \log\left(\bar{c}_1\right) + \left[\frac{1}{\bar{c}_1}\right] \left\{\left.\frac{\partial c_1(Z)}{\partial Z}\right|_{Z = \bar{Z}}\right\} (Z - \bar{Z})$$
$$+ \log\left(\bar{r}\right) + \left[\frac{1}{\bar{r}}\right] \left\{\left.\frac{\partial r(Z)}{\partial Z}\right|_{Z = \bar{Z}}\right\} (Z - \bar{Z})$$

$$- \log\left(e^{\theta_0+\theta_G\bar{G}+\theta_Z\bar{Z}}(1-\bar{c}_1)\bar{r}+1\right) - \frac{e^{\theta_0+\theta_G\bar{G}+\theta_Z\bar{Z}}(1-\bar{c}_1)\bar{r}}{e^{\theta_0+\theta_G\bar{G}+\theta_Z\bar{Z}}(1-\bar{c}_1)\bar{r}+1}\theta_G(G-\bar{G})$$

$$- \frac{e^{\theta_0+\theta_G\bar{G}+\theta_Z\bar{Z}}(1-\bar{c}_1)\bar{r}\theta_Z + e^{\theta_0+\theta_G\bar{G}+\theta_Z\bar{Z}}\left\{\frac{\partial(1-c_1(Z))r(Z)}{\partial Z}\Big|_{Z=\bar{Z}}\right\}}{e^{\theta_0+\theta_G\bar{G}+\theta_Z\bar{Z}}(1-\bar{c}_1)\bar{r}+1}(Z-\bar{Z})$$

**Suppose further that the covariate set $Z$ is centered** on the sample such that $\bar{Z}=0$. In this setting, we can rewrite the above expression as

$$\text{logit}\left(P(D^*=1|G,Z,S=1)\right) \approx \qquad \text{(SuppEq. 2.3)}$$

$$\theta_0 + \log(\bar{c}_1) + \log(\bar{r}) + \frac{e^{\theta_0+\theta_G\bar{G}}(1-\bar{c}_1)\bar{r}}{e^{\theta_0+\theta_G\bar{G}}(1-\bar{c}_1)\bar{r}+1}\theta_G\bar{G}$$

$$+ \left[\left[\frac{1}{\bar{c}_1}\right]\left\{\frac{\partial c_1(Z)}{\partial Z}\Big|_{Z=\bar{Z}}\right\} + \left[\frac{1}{\bar{r}}\right]\left\{\frac{\partial r(Z)}{\partial Z}\Big|_{Z=\bar{Z}}\right\} - \frac{e^{\theta_0+\theta_G\bar{G}}(1-\bar{c}_1)\bar{r}\theta_Z + e^{\theta_0+\theta_G\bar{G}}\left\{\frac{\partial(1-c_1(Z))r(Z)}{\partial Z}\Big|_{Z=\bar{Z}}\right\}}{e^{\theta_0+\theta_G\bar{G}}(1-\bar{c}_1)\bar{r}+1}\right] Z$$

$$+ \left[\frac{1}{e^{\theta_0+\theta_G\bar{G}}(1-\bar{c}_1)\bar{r}+1}\theta_G\right] G$$

We will consider two different cases for $G$. We will first suppose that $G$ represents a single genetic locus. Then, we will assume $G$ is any continuous predictor, but we will focus on the particular setting where $G$ is a polygenic risk score.

Case 1: $G$ is a SNP Suppose first that $G$ represents a single SNP (single nucleotide polymorphism) or genetic locus and that $G$ is coded 0/1/2, where 0 represents no copies of the minor allele, 1 represents one copy of the minor allele, and 2 represents two copies of the minor allele. We assume there are only two non-negligible alleles for the SNP of interest.

First, we replace $\bar{G}$ in the above derivation with $E(G|S=1)$. Let $MAF$ represent the minor allele frequency in the *population* of interest and $MAF^{(sam)}$ represent the minor allele frequency in the *sample*. Since these are quantities are averaged across $D$, $MAF$ and $MAF^{(sam)}$ may be different when sampling depends on $D$, which in turn depends on $G$. In practice, we usually don't expect $MAF$ to be too different from $MAF^{(sam)}$, so we will automatically replace $MAF^{(sam)}$ with $MAF$ in the approximation equations for $\theta_0^{(simple)}$ and $\theta_G^{(simple)}$ derived below.

Assuming that the two alleles for any given person are independent and assuming Hardy-Weinberg Equilibrium, we can approximate $\bar{G}$ roughly with

$$E(G|S=1) = \sum_{g=0,1,2} gP(G=g|S=1) = P(G=1|S=1) + 2*P(G=2|S=1)$$
$$\approx 2*MAF*(1-MAF) + 2*MAF^2 = 2*MAF$$

Substituting this expression for $\bar{G}$ in (SuppEq. 2.3), we can approximate $\theta_0^{(simple)}$ and $\theta_G^{(simple)}$ as follows:

Under Assumptions 1-4,

$$\theta_0^{(simple)} \approx \theta_0 + \log(\bar{c}_1) + \log(\bar{r}) - \log\left(e^{\theta_0+2\theta_G MAF}[1-\bar{c}_1]\bar{r}+1\right)$$
$$+ 2\theta_G MAF\left[\frac{e^{\theta_0+2\theta_G MAF}(1-\bar{c}_1)\bar{r}}{e^{\theta_0+2\theta_G MAF}(1-\bar{c}_1)\bar{r}+1}\right]$$
$$\theta_G^{(simple)} \approx \left[\frac{1}{e^{\theta_0+2\theta_G MAF}(1-\bar{c}_1)\bar{r}+1}\right]\theta_G$$

where

$\bar{c}_1 = \int f(D^* = 1|D = 1, X^\dagger, Z = 0, S = 1) f(X^\dagger|D = 1, Z = 0, S = 1) dX^\dagger = \text{Sensitivity}$
$\bar{r} = \frac{\int P(S=1|D=1,W^\dagger,Z=0) f(W^\dagger|D=1,Z=0) dW^\dagger}{\int P(S=1|D=0,W^\dagger,Z=0) f(W^\dagger|D=0,Z=0) dW^\dagger} = \text{Sampling Ratio with respect to } D$
and we assume $Z$ has been mean-centered, so $\bar{Z} = 0$.

Suppose we are in a setting in which $MAF^{(sam)}$ and $MAF$ may be somewhat different. In particular, suppose that sampling is somewhat strongly dependent on $D$ (so the sampling ratio is extreme) and $D$ is strongly related to $G$. In this setting, we may expect the sampling to impact the minor allele frequency in the sample compared to the entire population. In this setting, we may want to use $MAF^{(sam)}$ in the approximation equations for $\theta_0^{(simple)}$ and $\theta_G^{(simple)}$ in (SuppEq. 2.3) instead of $MAF$. In this case, we can approximate $MAF^{(sam)}$ in terms of $MAF$ as follows (replacing $Z$ with its mean, $\bar{Z} = 0$):

$$MAF^{(sam)} \approx MAF \frac{1 + (r-1)\left[MAF * P(D = 1|G = 2, Z = 0) + (1 - MAF)P(D = 1|G = 1, Z = 0)\right]}{1 + (r-1)P(D = 1)}$$

The derivation of this equation uses the decomposition,

$$\begin{aligned}
&MAF^{(sam)} = P(M = 1|S = 1) \\
&\approx \frac{\sum_{d,g} P(S = 1|D = d, W)P(D = d|G = g, Z = 0)P(G = g|M = 1, Z = 0)P(M = 1|Z = 0)f(W)}{P(S = 1)} \\
&= \frac{MAF}{P(S = 1)} MAF \sum_d \left[\int P(S = 1|D = d, W)f(W)dW\right] P(D = d|G = 2, Z = 0) \\
&+ \frac{MAF}{P(S = 1)}(1 - MAF) \sum_d \left[\int P(S = 1|D = d, W)f(W)dW\right] P(D = d|G = 1, Z = 0)
\end{aligned}$$

where $M$ is in indicator whether the first allele for a given subject is the minor allele, and we replace $P(M = 1|Z = 0)$ with $MAF$. We also assume that $P(G = g|Z = 0, M = 1) \approx P(G = g|M = 1)$.

The equation for $MAF^{(sam)}$ also depends on $P(D = 1)$. We can *roughly* express $P(D = 1)$ as follows

$$\begin{aligned}
P(D = 1) &= \int \sum_{g=0,1,2} P(D = 1|G = g, Z)f(G = g, Z)dZ \\
&\approx \sum_{g=0,1,2} P(D = 1|G = g, Z = \bar{Z} = 0)P(G = g|Z = 0) \\
&= \frac{e^{\theta_0}}{1 + e^{\theta_0}}(1 - MAF)^2 + \frac{e^{\theta_0+\theta_G}}{1 + e^{\theta_0+\theta_G}} 2 * MAF(1 - MAF) + \frac{e^{\theta_0+2\theta_G}}{1 + e^{\theta_0+2\theta_G}} MAF^2
\end{aligned}$$

Even when $MAF \neq MAF^{(sam)}$, we note that there is little impact of using $MAF$ over $MAF^{(sam)}$ in the bias expression derived above, so this difference may not matter much in practice. We note that $MAF = MAF^{(sam)}$ when $\theta_G = 0$ or $r = 1$.

Case 2: $G$ is a polygenic risk score Now, we assume that $G$ is a polygenic risk score (PRS) or some other continuous predictor. Suppose further that we have centered the polygenic risk score such that $\bar{G} = 0$ in the sampled datasets. In this case, we can directly use (SuppEq. 2.3) to obtain

Under Assumptions 1-4,

$$\theta_0^{(simple)} \approx \theta_0 + \log(\bar{c}_1) + \log(\bar{r}) - \log\left(e^{\theta_0}[1 - \bar{c}_1]\bar{r} + 1\right)$$

$$\theta_G^{(simple)} \approx \left[\frac{1}{e^{\theta_0}(1-\bar{c}_1)\bar{r}+1}\right]\theta_G$$

where
$\bar{c}_1 = \int f(D^* = 1|D = 1, X^\dagger, Z = 0, S = 1) f(X^\dagger|D = 1, Z = 0, S = 1) dX^\dagger$ = Sensitivity
$\bar{r} = \frac{\int P(S=1|D=1,W^\dagger,Z=0) f(W^\dagger|D=1,Z=0) dW^\dagger}{\int P(S=1|D=0,W^\dagger,Z=0) f(W^\dagger|D=0,Z=0) dW^\dagger}$ = Sampling Ratio with respect to $D$
and we assume $Z$ and the PRS have both been mean-centered.

##### General properties of bias

We are interested to see what will happen to $\theta_G^{(simple)}$ relative to $\theta_G$. We have that

$$\theta_G^{(simple)} \approx \left[\frac{1}{e^{\theta_0+\theta_G\bar{G}}(1-\bar{c}_1)\bar{r}+1}\right]\theta_G$$

This expression is exactly zero if $\bar{c}_1 = 1$, corresponding to perfect sensitivity. As $\bar{c}_1$ goes to 0, the parameter estimate is increasingly biased toward the null. The same is also true as $\bar{r}$ goes to infinity.

#### S2.2 Sensitivity analysis expressions

Suppose $\theta_G$ represents the association (log-odds ratio) between a particular genetic locus or PRS and the disease of interest. Rather than estimating $\theta_G$ directly, we most often fit a model for $D^*|G, Z, S = 1$ to obtain the estimate $\theta_G^{(simple)}$. We may be interested in exploring plausible values of $\theta_G$ given already known $\theta_G^{(simple)}$ and reasonable values of $\bar{r}$ and $\bar{c}_1$. Below, we describe how we can back out values for $\theta_G$ given $\bar{r}$, $\bar{c}_1$, $\theta_G^{(simple)}$, and either $\theta_0^{(simple)}$ or $\theta_0$.

Case 1: $G$ is a SNP Suppose first that we only know $\theta_G^{(simple)}$ but not $\theta_0^{(simple)}$ and that we have a plausible value for $\theta_0$ based on known population prevalence $P(D = 1)$ or $P(D = 1|G = 0, Z = \bar{Z})$. In practice, we can perform the following exploration using an interval of values for $\theta_0$. We can obtain a prediction for $\theta_G$ by *numerically solving*

$$\theta_G^{(simple)} \approx \left[\frac{1}{e^{\theta_0+2\theta_G MAF}(1-\bar{c}_1)\bar{r}+1}\right]\theta_G \qquad \text{(SuppEq. 2.4)}$$

for $\theta_G$. We note that this expression may have multiple solutions or no solutions for given values of $r$ and $\bar{c}_1$. Suppose instead that $\theta_G^{(simple)}$ and $\theta_0^{(simple)}$ are both available. In this case (assuming $\theta_G$ is not *too* far from 0, which may be reasonable when $G$ is a SNP), we approximate $\theta_0$ with

$$e^{\theta_0} \approx \frac{1}{r}\left[\frac{e^{\theta_0^{(simple)}}}{\bar{c}_1 - e^{\theta_0^{(simple)}}(1-\bar{c}_1)}\right]$$

We obtain the above expression by solving the bias expression for $\theta_0^{(simple)}$ and setting $\theta_G = 0$. We can plug the above approximation into expression (SuppEq. 2.4). Based on this and assuming $\theta_G$ is somewhat near zero, we can obtain a plausible value for $\theta_G$ by solving the following:

$$\theta_G^{(simple)} \approx \frac{\bar{c}_1 - e^{\theta_0^{(simple)}}(1-\bar{c}_1)}{\bar{c}_1 - e^{\theta_0^{(simple)}}(1-\bar{c}_1)\left[1-e^{2\theta_G MAF}\right]}\theta_G \qquad \text{(SuppEq. 2.5)}$$

for $\theta_G$. We note that this expression does not depend on $\bar{r}$. This is due to the approximations made and the fact that both $\theta_G^{(simple)}$ and $\theta_0^{(simple)}$ are provided.

Case 2: $G$ is a PRS This case is simpler, where we can directly express $\theta_G$ as

$$\theta_G \approx \theta_G^{(simple)}\left[e^{\theta_0}(1-\bar{c}_1)\bar{r}+1\right] \qquad \text{(SuppEq. 2.6)}$$

and replace $\theta_0$ with an assumed value. When $\theta_0^{(simple)}$ is also available, we can alternatively use the expression

$$\theta_G \approx \theta_G^{(simple)}\left[\frac{\bar{c}_1}{\bar{c}_1 - e^{\theta_0^{(simple)}}(1-\bar{c}_1)}\right] \qquad \text{(SuppEq. 2.7)}$$

to predict $\theta_G$ given $\bar{c}_1$ and $\theta^{(simple)}$

#### S3 Bias and sensitivity analysis under imperfect specificity

##### S3.1 Bias expressions

We will first assume that we fit analysis model 1 as in SuppEq. 1.2 and allow specificity to be less than 1. Define

$$c_0(Z;\alpha) = \int f(D^* = 1|D = 0, Y^\dagger, Z, S = 1) f(Y^\dagger|D = 0, Z, S = 1) dY^\dagger$$

We can view $c_0(Z)$ as one minus the specificity of $D^*$ for $D$ in the sampled subjects averaged across the distribution of $Y^\dagger$. $c_1(Z)$ and $r(Z)$ are defined as before.

Suppose we use logistic regression to model $D^*|G, Z, S = 1$ as in (SuppEq. 1.1) analysis model 1. We have

$$\begin{aligned}
&\text{logit}\,(P(D^* = 1|G, Z, S = 1)) \\
&= \log\left(\frac{P(D = 1|G, Z)c_1(Z)r(Z) + P(D = 0|G, Z)c_0(Z)}{P(D = 1|G, Z)(1 - c_1(Z))r(Z) + P(D = 0|G, Z)(1 - c_0(Z))}\right) \\
&= \log\left(\frac{e^{\theta_0+\theta_G G+\theta_Z Z}c_1(Z)r(Z) + c_0(Z)}{e^{\theta_0+\theta_G G+\theta_Z Z}(1 - c_1(Z))r(Z) + (1 - c_0(Z))}\right) \\
&= \log\left(\frac{c_0(Z)}{1 - c_0(Z)}\right) + \log\left(e^{\theta_0+\theta_G G+\theta_Z Z}\frac{c_1(Z)r(Z)}{c_0(Z)} + 1\right) - \log\left(e^{\theta_0+\theta_G G+\theta_Z Z}\frac{(1 - c_1(Z))r(Z)}{1 - c_0(Z)} + 1\right)
\end{aligned}$$

We now approximate the above expression using a first order Taylor Series approximation with respect to $Z$ and $G$, where $\bar{Z}$ represents the mean of $Z$ and $\bar{G}$ represents the mean of $G$ in the sample. Let $\bar{c}_1 = c_1(\bar{Z})$, $\bar{c}_0 = c_0(\bar{Z})$, and $\bar{r} = r(\bar{Z})$. We can write

$$\begin{aligned}
&\text{logit}\,(P(D^* = 1|G, Z, S = 1)) \approx \log\left(\frac{\bar{c}_0}{1 - \bar{c}_0}\right) + \left\{\frac{1}{\bar{c}_0} + \frac{1}{1 - \bar{c}_0}\right\}\left\{\frac{\partial c_0(Z)}{\partial Z}\Big|_{Z=\bar{Z}}\right\}(Z - \bar{Z}) \\
&+ \log\left(e^{\theta_0+\theta_G\bar{G}+\theta_Z\bar{Z}}\frac{\bar{c}_1\bar{r}}{\bar{c}_0} + 1\right) + \frac{e^{\theta_0+\theta_G\bar{G}+\theta_Z\bar{Z}}\frac{\bar{c}_1\bar{r}}{\bar{c}_0}}{e^{\theta_0+\theta_G\bar{G}+\theta_Z\bar{Z}}\frac{\bar{c}_1\bar{r}}{\bar{c}_0} + 1}\theta_G(G - \bar{G}) \\
&+ \frac{e^{\theta_0+\theta_G\bar{G}+\theta_Z\bar{Z}}\frac{\bar{c}_1\bar{r}}{\bar{c}_0}\theta_Z + e^{\theta_0+\theta_G\bar{G}+\theta_Z\bar{Z}}\left\{\frac{\partial\frac{c_1(Z)r(Z)}{c_0(Z)}}{\partial Z}\Big|_{Z=\bar{Z}}\right\}}{e^{\theta_0+\theta_G\bar{G}+\theta_Z\bar{Z}}\frac{\bar{c}_1\bar{r}}{\bar{c}_0} + 1}(Z - \bar{Z}) \\
&- \log\left(e^{\theta_0+\theta_G\bar{G}+\theta_Z\bar{Z}}\frac{(1 - \bar{c}_1)\bar{r}}{1 - \bar{c}_0} + 1\right) - \frac{e^{\theta_0+\theta_G\bar{G}+\theta_Z\bar{Z}}\frac{(1-\bar{c}_1)\bar{r}}{1-\bar{c}_0}}{e^{\theta_0+\theta_G\bar{G}+\theta_Z\bar{Z}}\frac{(1-\bar{c}_1)\bar{r}}{1-\bar{c}_0} + 1}\theta_G(G - \bar{G}) \\
&- \frac{e^{\theta_0+\theta_G\bar{G}+\theta_Z\bar{Z}}\frac{(1-\bar{c}_1)\bar{r}}{1-\bar{c}_0}\theta_Z + e^{\theta_0+\theta_G\bar{G}+\theta_Z\bar{Z}}\left\{\frac{\partial\frac{(1-c_1(Z))r(Z)}{1-c_0(Z)}}{\partial Z}\Big|_{Z=\bar{Z}}\right\}}{e^{\theta_0+\theta_G\bar{G}+\theta_Z\bar{Z}}\frac{(1-\bar{c}_1)\bar{r}}{1-\bar{c}_0} + 1}(Z - \bar{Z})
\end{aligned}$$

**Suppose further that the covariate set $Z$ is centered** on the sample such that $\bar{Z} = 0$. In this setting, we can rewrite the above expression as

$$\begin{aligned}
&\text{logit}\,(P(D^* = 1|G, Z, S = 1)) \approx \qquad \text{(SuppEq. 3.8)} \\
&\log\left(\frac{e^{\theta_0+\theta_G\bar{G}}\bar{c}_1\bar{r} + \bar{c}_0}{e^{\theta_0+\theta_G\bar{G}}[1 - \bar{c}_1]\bar{r} + [1 - \bar{c}_0]}\right) - \frac{e^{\theta_0+\theta_G\bar{G}}\frac{\bar{c}_1\bar{r}}{\bar{c}_0}}{e^{\theta_0+\theta_G\bar{G}}\frac{\bar{c}_1\bar{r}}{\bar{c}_0} + 1}\theta_G\bar{G} + \frac{e^{\theta_0+\theta_G\bar{G}}\frac{(1-\bar{c}_1)\bar{r}}{1-\bar{c}_0}}{e^{\theta_0+\theta_G\bar{G}}\frac{(1-\bar{c}_1)\bar{r}}{1-\bar{c}_0} + 1}\theta_G\bar{G} \\
&+ \Bigg[\left\{\frac{1}{\bar{c}_0} + \frac{1}{1 - \bar{c}_0}\right\}\left\{\frac{\partial c_0(Z)}{\partial Z}\Big|_{Z=0}\right\} - \frac{e^{\theta_0+\theta_G\bar{G}}\frac{(1-\bar{c}_1)\bar{r}}{1-\bar{c}_0}\theta_Z + e^{\theta_0+\theta_G\bar{G}}\left\{\frac{\partial\frac{(1-c_1(Z))r(Z)}{1-c_0(Z)}}{\partial Z}\Big|_{Z=0}\right\}}{e^{\theta_0+\theta_G\bar{G}}\frac{(1-\bar{c}_1)\bar{r}}{1-\bar{c}_0} + 1}
\end{aligned}$$

$$+\frac{e^{\theta_0+\theta_G\bar{G}}\frac{\bar{c}_1\bar{r}}{\bar{c}_0}\theta_Z + e^{\theta_0+\theta_G\bar{G}}\left\{\frac{\partial\frac{c_1(Z)r(Z)}{c_0(Z)}}{\partial Z}\Bigg|_{Z=0}\right\}}{e^{\theta_0+\theta_G\bar{G}}\frac{\bar{c}_1\bar{r}}{\bar{c}_0}+1}\Bigg] Z + \left[\frac{e^{\theta_0+\theta_G\bar{G}}\frac{\bar{c}_1\bar{r}}{\bar{c}_0}}{e^{\theta_0+\theta_G\bar{G}}\frac{\bar{c}_1\bar{r}}{\bar{c}_0}+1}\theta_G - \frac{e^{\theta_0+\theta_G\bar{G}}\frac{(1-\bar{c}_1)\bar{r}}{1-\bar{c}_0}}{e^{\theta_0+\theta_G\bar{G}}\frac{(1-\bar{c}_1)\bar{r}}{1-\bar{c}_0}+1}\theta_G\right] G$$

As before, we will consider two different cases for $G$. We will first suppose that $G$ represents a single genetic locus. Then, we will assume $G$ is any continuous predictor, but we will focus on the particular setting where $G$ is a polygenic risk score.

Case 1: Bias expressions assuming $G$ is a SNP Suppose first that $G$ represents a single SNP (single nucleotide polymorphism) or genetic locus and that $G$ is coded 0/1/2, where 0 represents no copies of the minor allele, 1 represents one copy of the minor allele, and 2 represents two copies of the minor allele. We assume there are only two non-negligible alleles for the SNP of interest. Let $MAF$ and $MAF^{(sam)}$ be as in **Section S2**. Substituting this expression for $\bar{G}$ in (SuppEq. 3.8), we can approximate $\theta_0^{(simple)}$ and $\theta_G^{(simple)}$ as follows:

Under Assumptions 1-4,

$$\theta_0^{(simple)} \approx \log\left(\frac{e^{\theta_0+2\theta_G MAF}\bar{c}_1\bar{r}+\bar{c}_0}{e^{\theta_0+2\theta_G MAF}[1-\bar{c}_1]\bar{r}+[1-\bar{c}_0]}\right) - 2\theta_G MAF\left[\frac{e^{\theta_0+2\theta_G MAF}\frac{\bar{c}_1\bar{r}}{\bar{c}_0}}{e^{\theta_0+2\theta_G MAF}\frac{\bar{c}_1\bar{r}}{\bar{c}_0}+1} - \frac{e^{\theta_0+2\theta_G MAF}\frac{(1-\bar{c}_1)\bar{r}}{1-\bar{c}_0}}{e^{\theta_0+2\theta_G MAF}\frac{(1-\bar{c}_1)\bar{r}}{1-\bar{c}_0}+1}\right]$$

$$\theta_G^{(simple)} \approx \left[\frac{e^{\theta_0+2\theta_G MAF}\frac{\bar{c}_1\bar{r}}{\bar{c}_0}}{e^{\theta_0+2\theta_G MAF}\frac{\bar{c}_1\bar{r}}{\bar{c}_0}+1} - \frac{e^{\theta_0+2\theta_G MAF}\frac{(1-\bar{c}_1)\bar{r}}{1-\bar{c}_0}}{e^{\theta_0+2\theta_G MAF}\frac{(1-\bar{c}_1)\bar{r}}{1-\bar{c}_0}+1}\right]\theta_G \qquad \text{(SuppEq. 3.9)}$$

where
$\bar{c}_1 = \int f(D^*=1|D=1,X^\dagger,Z=0,S=1)f(X^\dagger|D=1,Z=0,S=1)dX^\dagger$ = Sensitivity
$\bar{c}_0 = \int f(D^*=1|D=0,Y^\dagger,Z=0,S=1)f(Y^\dagger|D=0,Z=0,S=1)dY^\dagger$ = 1- Specificity
$\bar{r} = \frac{\int P(S=1|D=1,W^\dagger,Z=0)f(W^\dagger|D=1,Z=0)dW^\dagger}{\int P(S=1|D=0,W^\dagger,Z=0)f(W^\dagger|D=0,Z=0)dW^\dagger}$ = Sampling Ratio with respect to $D$
and we assume $Z$ has been mean-centered, so $\bar{Z}=0$.

Case 2: Bias expressions assuming $G$ is a polygenic risk score Now, we assume that $G$ is a polygenic risk score (PRS) or some other continuous predictor. Suppose further that we have centered the polygenic risk score such that $\bar{G}=0$ in the sampled datasets. In this case, we can directly use (SuppEq. 3.8) to obtain

Under Assumptions 1-4,

$$\theta_0^{(simple)} \approx \log\left(\frac{e^{\theta_0}\bar{c}_1\bar{r}+\bar{c}_0}{e^{\theta_0}[1-\bar{c}_1]\bar{r}+[1-\bar{c}_0]}\right)$$

$$\theta_G^{(simple)} \approx \left[\frac{e^{\theta_0}\frac{\bar{c}_1\bar{r}}{\bar{c}_0}}{e^{\theta_0}\frac{\bar{c}_1\bar{r}}{\bar{c}_0}+1} - \frac{e^{\theta_0}\frac{(1-\bar{c}_1)\bar{r}}{1-\bar{c}_0}}{e^{\theta_0}\frac{(1-\bar{c}_1)\bar{r}}{1-\bar{c}_0}+1}\right]\theta_G \qquad \text{(SuppEq. 3.10)}$$

where
$\bar{c}_1 = \int f(D^*=1|D=1,X^\dagger,Z=0,S=1)f(X^\dagger|D=1,Z=0,S=1)dX^\dagger$ = Sensitivity
$\bar{c}_0 = \int f(D^*=1|D=0,Y^\dagger,Z=0,S=1)f(Y^\dagger|D=0,Z=0,S=1)dY^\dagger$ = 1- Specificity
$\bar{r} = \frac{\int P(S=1|D=1,W^\dagger,Z=0)f(W^\dagger|D=1,Z=0)dW^\dagger}{\int P(S=1|D=0,W^\dagger,Z=0)f(W^\dagger|D=0,Z=0)dW^\dagger}$ = Sampling Ratio with respect to $D$
and we assume $Z$ and the PRS have both been mean-centered.

General properties of bias We are interested to see what will happen to $\theta_G^{(simple)}$ relative to

$\theta_G$ in various settings. First, we rewrite the above expression as follows:

$$\theta_G^{(simple)} \approx \left[ \frac{\frac{\bar{c}_1}{\bar{c}_0} - \frac{(1-\bar{c}_1)}{1-\bar{c}_0}}{\left[e^{\theta_0+\theta_G\bar{G}}\frac{\bar{c}_1\bar{r}}{\bar{c}_0}+1\right]\left[e^{\theta_0+\theta_G\bar{G}}\frac{(1-\bar{c}_1)\bar{r}}{1-\bar{c}_0}+1\right]} \right] \theta_G e^{\theta_0+\theta_G\bar{G}}\bar{r}$$

This expression is exactly zero if $\bar{c}_1 = \bar{c}_0$. This corresponds to the setting where the probability of diagnosing a subject with a disease is equal among diseased and non-diseased subjects. This is a setting we do not expect to encounter in practice. We will therefore assume $\bar{c}_1 > \bar{c}_0$.

A particularly concerning setting is when $\theta_G^{(simple)}$ is in the opposite direction as $\theta_G$. This occurs when $\bar{c}_1(1-\bar{c}_0) < (1-\bar{c}_1)\bar{c}_0$ or equivalently when $sens * spec < (1 - sens) * (1 - spec)$ where *sens* represents the sensitivity and *spec* represents the specificity. Suppose that we have either a specificity or a sensitivity of 1. In that case, this expression is never satisfied, and we will never have direction switching. Suppose, however, that both are not equal to 1. In this case, it is possible to have direction switching. While we do not expect specificity to be below 0.5, it is theoretically possible to have very low sensitivity for observing disease status for EHR-based observations. In settings in which sensitivity is below 0.5, we can have direction switching, which may be particular troubling when the goal is to study the direction and strength of an association.

Suppose that $\bar{c}_1 = 1 - \bar{c}_0$. This is the setting of non-differential outcome misclassification. In this setting, we could have direction switching if $sens^2 < (1 - sens)^2$, so if *sens* is less than 0.5.

We are also interested in determining when $\theta_G^{(simple)}$ will be biased toward or away from the null value of zero. It is difficult to specify general rules, and this should be evaluated on a case-by-case basis for plausible values of $\bar{r}$, $\bar{c}_1$, and $\bar{c}_0$. To the extent possible, **Table S1** provides intuition regarding the bias for $\theta_G^{(simple)}$ under perfect and imperfect specificity.

##### S3.2 Sensitivity analysis expressions

Case 1: $G$ is a SNP Suppose that $G$ is a SNP as described in **Section S2**. Suppose first that we only know $\theta_G^{(simple)}$ but not $\theta_0^{(simple)}$. This would be the case if we were exploring bias using published GWAS results or using GWAS pipelines that do not routinely save $\theta_0^{(simple)}$. Suppose instead that we have a plausible value for $\theta_0$ based on known population prevalence $P(D=1)$ or $P(D=1|G=0,Z=\bar{Z})$. In practice, we can perform the following exploration using an interval of values for $\theta_0$. We can obtain a prediction for $\theta_G$ by *numerically solving*

$$\theta_G^{(simple)} \approx \left[ \frac{e^{\theta_0+2\theta_G MAF \frac{\bar{c}_1 \bar{r}}{\bar{c}_0}}}{e^{\theta_0+2\theta_G MAF \frac{\bar{c}_1 \bar{r}}{\bar{c}_0}}+1} - \frac{e^{\theta_0+2\theta_G MAF \frac{(1-\bar{c}_1)\bar{r}}{1-\bar{c}_0}}}{e^{\theta_0+2\theta_G MAF \frac{(1-\bar{c}_1)\bar{r}}{1-\bar{c}_0}}+1} \right] \theta_G$$

for $\theta_G$. We note that this expression may have multiple solutions or no solutions for given values of $r$, $\bar{c}_1$ and $\bar{c}_0$. When there are multiple solutions, we will choose the solution closest to $\theta_G^{(simple)}$. Simulations exploring the compatibility of these expressions with simulated data can be found later on in **Section S6** and in the main paper.

Suppose instead that $\theta_G^{(simple)}$ and $\theta_0^{(simple)}$ are both available. In this case (assuming $\theta_G$ is not *too* far from 0, which may be reasonable when $G$ is a SNP), we approximate $\theta_0$ with

$$e^{\theta_0} \approx \frac{1}{\bar{r}} \left[ \frac{e^{\theta_0^{(simple)}}(1-\bar{c}_0) - \bar{c}_0}{\bar{c}_1 - e^{\theta_0^{(simple)}}(1-\bar{c}_1)} \right]$$

We obtain the above expression by solving the bias expression for $\theta_0^{(simple)}$ setting $\theta_G = 0$. Based on this and assuming $\theta_G$ is somewhat near zero, we can obtain a plausible value for $\theta_G$ by solving the following:

$$\theta_G^{(simple)} \approx \left[ \frac{\left[\frac{e^{\theta_0^{(simple)}}\frac{(1-\bar{c}_0)}{\bar{c}_0}-1}{1-e^{\theta_0^{(simple)}}\frac{(1-\bar{c}_1)}{\bar{c}_1}}\right] e^{2\theta_G MAF}}{\left[\frac{e^{\theta_0^{(simple)}}\frac{(1-\bar{c}_0)}{\bar{c}_0}-1}{1-e^{\theta_0^{(simple)}}\frac{(1-\bar{c}_1)}{\bar{c}_1}}\right] e^{2\theta_G MAF}+1} - \frac{\left[\frac{e^{\theta_0^{(simple)}}-\frac{\bar{c}_0}{1-\bar{c}_0}}{\frac{\bar{c}_1}{1-\bar{c}_1}-e^{\theta_0^{(simple)}}}\right] e^{2\theta_G MAF}}{\left[\frac{e^{\theta_0^{(simple)}}-\frac{\bar{c}_0}{1-\bar{c}_0}}{\frac{\bar{c}_1}{1-\bar{c}_1}-e^{\theta_0^{(simple)}}}\right] e^{2\theta_G MAF}+1} \right] \theta_G$$

for $\theta_G$. We note that this expression does not depend on $\bar{r}$. This is due to the approximations made and the fact that both $\theta_G^{(simple)}$ and $\theta_0^{(simple)}$ are provided.

Case 2: $G$ is a PRS This case is simpler, where we can directly express $\theta_G$ as

$$\theta_G \approx \theta_G^{(simple)} \left[ \frac{e^{\theta_0}\frac{\bar{c}_1 \bar{r}}{\bar{c}_0}}{e^{\theta_0}\frac{\bar{c}_1 \bar{r}}{\bar{c}_0}+1} - \frac{e^{\theta_0}\frac{(1-\bar{c}_1)\bar{r}}{1-\bar{c}_0}}{e^{\theta_0}\frac{(1-\bar{c}_1)\bar{r}}{1-\bar{c}_0}+1} \right]^{-1}$$

and replace $\theta_0$ with an assumed value. When $\theta_0^{(simple)}$ is also available, we can alternatively use the expression

$$\theta_G \approx \theta_G^{(simple)} \left[ \frac{\left[\frac{e^{\theta_0^{(simple)}}\frac{(1-\bar{c}_0)}{\bar{c}_0}-1}{1-e^{\theta_0^{(simple)}}\frac{(1-\bar{c}_1)}{\bar{c}_1}}\right]}{\left[\frac{e^{\theta_0^{(simple)}}\frac{(1-\bar{c}_0)}{\bar{c}_0}-1}{1-e^{\theta_0^{(simple)}}\frac{(1-\bar{c}_1)}{\bar{c}_1}}\right]+1} - \frac{\left[\frac{e^{\theta_0^{(simple)}}-\frac{\bar{c}_0}{1-\bar{c}_0}}{\frac{\bar{c}_1}{1-\bar{c}_1}-e^{\theta_0^{(simple)}}}\right]}{\left[\frac{e^{\theta_0^{(simple)}}-\frac{\bar{c}_0}{1-\bar{c}_0}}{\frac{\bar{c}_1}{1-\bar{c}_1}-e^{\theta_0^{(simple)}}}\right]+1} \right]^{-1}$$

to predict $\theta_G$ given $\bar{c}_1$, $\bar{c}_0$, and $\theta^{(simple)}$

#### S4 Second stage sampling based only on observed phenotypes

In **Section S2**, we considered sampling into the large observational dataset (such as the EHR) and allowed sampling to depend on patient characteristics $W$ and on the *underlying disease status* $D$. In practice, researchers often obtain their analytical sample from the larger dataset (e.g. all subjects with available EHR information) based on the *observed* disease status, $D^*$ (e.g whether subjects are diagnosed with diabetes). We can view this is case-control sampling with respect to the outcome of the *analysis model*, $D^*$.

Here, we will make a distinction between the mechanism in which we select people from the target population into the large dataset (e.g. the set of patients visiting a particular hospital) and the mechanism in which we sample patients from this larger dataset into our analytical sample (e.g. a subset of subjects visiting a particular hospital). We will assume the case-control sampling is based on the disease of interest. If the case-control sampling is based on the observed status for a *different* disease, additional thinking will be required.

We define $S_1$ to be an indicator for sampling into the large dataset (e.g into the hospital) and $S_2$ to represent sampling into our analytical sample. $S_1 = 0$ automatically implies $S_2 = 0$. $S_1$ is sometimes defined by non-probability sampling in which we do not know the true sampling mechanism, but we will place some assumptions on the structure of this model as before (that it depends on covariates $W$ and possibly on $D$). We assume $S_2$ depends only on $D^*$ given $S_1 = 1$. **Figure S1** shows the assumed data structure.

We are interested in exploring the bias of parameters in the *analysis model*, but this time the model we are actually fitting is for $D^*|G, Z, S_2 = 1$. We can show that

$$f(D^*|G, Z, S_2 = 1) = \frac{P(S_2 = 1|D^*, S_1 = 1)f(D^*|S_1, G, Z)P(S_1 = 1|G, Z)}{P(S_2 = 1|G, Z)}$$

and

$$\text{logit}(P(D^* = 1|G, Z, S_2 = 1)) = \log(r^{cc}) + \log\left(\frac{P(D^* = 1|S_1, G, Z)}{P(D^* = 0|S_1, G, Z)}\right)$$

where $r^{cc} = \frac{P(S_2=1|D^*=1,S_1=1)}{P(S_2=1|D^*=0,S_1=1)}$ is the case-control sampling ratio among the large dataset (with $S_1 = 1$). Since $r^{cc}$ is a constant with respect to $\theta_G$, it will not impact the derivation of the approximated formula for $\theta_G^{(simple)}$. The approximation for $\theta_0^{(simple)}$ will be the same except with an additional offset term $\log(r^{cc})$. Therefore, a final stage of case-control sampling dependent on $D^*$ will not induce additional bias in the estimation of $\theta_G^{(simple)}$ beyond bias due to the sampling mechanism related to $S_1$ and the observation/misclassification mechanisms for $D$.

#### S5 Secondary analysis of case-control data and sampling on other diseases

**Suppose that we have no misclassification of the outcome data.** In this section, we will explore the impact of different sampling mechanisms and how they fit into our independence assumptions and work done in the literature for secondary analyses for case-control data.

Suppose we have the following sampling mechanism: $P(S = 1|W, D; \phi)$, where we again assume that $G$ is not *independently* related to the sampling mechanism. However, $W$ can be marginally related to $D$, $G$, and $Z$. Define $W^\dagger$ as before. Using properties of logistic regression, can write the distribution of $D$ in the sampled subjects as

$$\begin{aligned}\text{logit}(P(D = 1|G, Z, S = 1)) &= \theta_0 + \theta_G G + \theta_Z Z + \log\left[\frac{\int P(S = 1|D = 1, W) f(W^\dagger|D = 1, G, Z) dW^\dagger}{\int P(S = 1|D = 0, W) f(W^\dagger|D = 0, G, Z) dW^\dagger}\right] \\ &= \theta_0 + \theta_G G + \theta_Z Z + \log\left[r(Z, G)\right]\end{aligned}$$

##### S5.1 $W$ included in $Z$

Suppose first that $W^\dagger$ is empty, so we include all additional factors driving sampling in $Z$. In this case, we have

$$\text{logit}(P(D = 1|G, Z, S = 1)) = \theta_0 + \theta_G G + \theta_Z Z + \log\left[\frac{P(S = 1|D = 1, W)}{P(S = 1|D = 0, W)}\right]$$

Suppose first that $D$ is *not independently* related to the sampling mechanism given $W \in Z$. In this setting, we will not expect any bias in estimating $\theta_G$ because $r(Z, G) = 1$ in that setting.

We are more interested in the setting in which $D$ is independently related to sampling given $W$, so $r(Z, G)$ is not uniformly equal to 1. Suppose we fit a logistic regression model incorrectly adjusting *linearly* for $G$ and $Z$ (which includes $W$). Adjustment for $W$ in the analysis model is one strategy for handling sampling dependent on another disease status in the case-control sampling literature. This can reduce bias in some cases, but not always [1, 2]. This strategy may not completely remove the bias in $\theta_G^{(simple)}$, but it may often have improved performance over ignoring sampling dependence on $W$. We note that our bias expressions are derived using first order Taylor series approximations. A first order Taylor series approximation of the above equation is

$$\begin{aligned}\text{logit}(P(D = 1|G, Z, S = 1)) &\approx \theta_0 + \theta_G G + \theta_Z Z + \log(P(S = 1|D = 1, \bar{W})) + \frac{\frac{\partial P(S=1|D=1,\bar{W})}{\partial W}}{P(S = 1|D = 1, \bar{W})} \\ &- \log(P(S = 1|D = 0, \bar{W})) - \frac{\frac{\partial P(S=1|D=0,\bar{W})}{\partial W}}{P(S = 1|D = 0, \bar{W})} \\ &= \left[\theta_0 + \log(P(S = 1|D = 1, \bar{W})) - \log(P(S = 1|D = 0, \bar{W}))\right] + \theta_G G \\ &+ \left[\theta_Z Z + \frac{\frac{\partial P(S=1|D=1,\bar{W})}{\partial W}}{P(S = 1|D = 1, \bar{W})} - \frac{\frac{\partial P(S=1|D=0,\bar{W})}{\partial W}}{P(S = 1|D = 0, \bar{W})}\right]\end{aligned}$$

In the first order Taylor series approximation, the coefficient for $G$ does not change. This suggests that $\theta_G^{(simple)}$ might be a reasonable approximation for $\theta_G$. While existing literature suggests that this may not always be a good approximation, this analysis approach still gives us some rough idea of the ballpark of $\theta_G$, which is the goal of this paper. We may expect the bias of $\theta_G^{(simple)}$ to be less of a concern when $W$ is independent of $G$ given $Z$ and $D$, since the incorrect modeling of $W$ may more greatly impact estimated $\theta_G^{(simple)}$ when $W$ and $G$ are associated.

In all settings in which $W$ is included in $Z$ explored in this paper, we do not make any assumptions on the relationship between $W$ and $G$ in this paper (assumption 4 in **Section S2** is trivial since $W^\dagger$ is empty in that case). Instead, we rely on the above results and focus instead on the impact of the dependence between sampling and $D$, adjusting for $W$. This strategy is expected to have good performance except when $W$ is strongly related to $G$ given $D$ and $Z$. In this setting, we may have appreciably biased estimates of $\theta_G^{(simple)}$ even after adjusting for $W$.

##### S5.2 $W$ not included in $Z$

More commonly, however, we may not adjust for all factors in $W$ in our simple analysis model. Consider the setting where $W$ represents a single secondary disease, $D'$, which may be related $G$ and/or $D$ given $Z$. In this case, we can rewrite the overall sampling mechanism as $P(S = 1|D, D')$ and express the correct model as

$$\begin{aligned}\text{logit}(P(D=1|G,Z,S=1)) &= \theta_0 + \theta_G G + \theta_Z Z + \log\left[\frac{\int P(S=1|D=1,D')f(D'|D=1,G,Z)dD'}{\int P(S=1|D=0,D')f(D'|D=0,G,Z)dD'}\right] \\ &= \theta_0 + \theta_G G + \theta_Z Z + \log\left[r(Z,G)\right]\end{aligned}$$

If we assume that $D' = W$ is independent of $G$ given $Z$ and $D$ (assumption 4 in **Section S2**), then we can rewrite the above expression as

$$\begin{aligned}\text{logit}(P(D=1|G,Z,S=1)) &= \theta_0 + \theta_G G + \theta_Z Z + \log\left[\frac{\int P(S=1|D=1,D')f(D'|D=1,Z)dD'}{\int P(S=1|D=0,D')f(D'|D=0,Z)dD'}\right] \\ &= \theta_0 + \theta_G G + \theta_Z Z + \log\left[r(Z)\right]\end{aligned}$$

Under logic from **Section S5.1**, we do not expect too much bias in estimating $\theta_G$ in this case, although there may still be some [3]. We note that this independence assumption conditions on $D$. Therefore, we will be okay if the relationship between $D'$ and $G$ is through $D$.

Suppose, however, that we have a disease $D'$ that is independently related to $G$ given $D$. An example of this would be the setting of pleiotropy. This may also be the case if $G$ represents a PRS. In this case, the offset term $r$ is a function of $G$, and failure to account for the missingness mechanism by fitting the simple analysis model can result in biased estimates. We consider this setting in more detail in **Section S9**.

#### S6 Comparison of approximations with simulated values

##### S6.1 Evaluating expressions for bias

The usefulness of the expressions in **Sections S3.1 and S2.1** relies on the accuracy of the Taylor series approximations used to derive them. In this section, we compare the approximated $\theta^{(simple)}$ with values obtained through simulation. We consider the setting where $G$ is a mean-centered polygenic risk score or some other continuous covariate and the setting where $G$ represents a single genetic locus.

Below, we provide details regarding how the data were simulated in each case. Briefly, we consider 96 different combinations of $\bar{r}$, $\bar{c}_1$, $\bar{c}_0$, and $\theta_0$. For each combination, we obtain 100 (Case 1) or 50 (Case 2) simulated datasets and fit the simple analysis model to each dataset. We then plot the approximated $\theta_G^{(simple)}$ from **Section S3.1** if $\bar{c}_0 > 0$ or **Section S2.1** if $\bar{c}_0 = 0$ against the median estimated $\theta_G^{(simple)}$ across the simulations for each of the 96 simulation settings. Results are shown in **Figure S2**.

We can see that there is generally excellent correspondence between the simulation-estimated $\theta^{(simple)}$ and the values obtained using the approximations in **Sections S3.1 and S2.1**, particularly for $\theta_0^{(simple)}$. An improved approximation of $\theta_G^{(simple)}$ may be obtained by taking second order Taylor Series approximations to obtain the approximation formulas. An exception is the setting in which we have very low sensitivity and low population prevalence of disease. In this setting, we sometimes see deviation between the approximation-predicted $\theta_G^{(simple)}$ and the estimated value, particularly when $\bar{r}$ is not very large (e.g. less than 2). In this setting, we may have very small numbers of subjects with $D^* = 1$, resulting in numerical challenges with fitting the logistic regression. These deviations result from difficulty in obtaining the simulation estimates rather than deficiencies of the approximation formula.

Simulating data when $G$ is a SNP First, we consider the setting where $G$ represents a SNP, coded 0/1/2 to reflect the number of minor alleles. We perform the following for each of several parameter values. We simulate 100 datasets of 50,000 subjects each under the conceptual model presented in (SuppEq. 1.1). For each dataset, we simulate covariate $G$ from a multinomial distribution with probabilities $[(1-MAF)^2, 2*MAF(1-MAF), MAF^2]$ for $[0, 1, 2]$ respectively with $MAF$ equal to 0.2. For each subject, we generate $D$ using $P(D = 1|G) = \text{expit}(\theta_0 + 0.5G)$. Here, there are no additional covariates $Z$ included in the model. We can view this as setting $Z = 0$ for all subjects. We then generate $S$ using $P(S = 1|D) = 0.1(1 - D) + 0.1\bar{r}D$. For the subjects with $S = 1$ and $D = 0$, we generate $D^*$ using $P(D^* = 1|S = 1, D = 0) = \bar{c}_0$. For subjects with $S = 1$ and $D = 1$, we generate $D^*$ using $P(D^* = 1|S = 1, D = 1) = \bar{c}_1$. We simulate these datasets under all combinations of $\bar{c}_1 = (0.1, 0.4, 0.7, 0.9)$, $\bar{c}_0 = (0, 0.05)$, $\bar{r} = (1, 2, 5, 10)$, and $\theta_0 = \text{logit}(0.01, 0.05, 0.10, 0.25)$. This results in a total of 128 simulation settings.

Simulating data when $G$ is a PRS Now, we consider the simpler setting where $G$ is a mean-centered PRS or some other mean-centered continuous variable. Note that we are still assuming that $G$ is independent of both the sampling and underreporting mechanisms given $D$. We simulate data as in Case 1 expect this time we simulate $G \sim N(0, 1)$ and use 50 simulated datasets.

#### S6.2 Evaluating proposed sensitivity analysis

In **Sections S3.2 and S2.2**, we derived equations we can use in a sensitivity analysis to explore plausible values of $\theta_G$ given $\theta_G^{(simple)}$ and either $\theta_0$ or estimated $\theta_0^{(simple)}$. In this section, we explore the concordance of the predicted values of $\theta_G$ with true values via simulation. We will focus on the setting where $G$ is a SNP as this setting is more complicated.

We simulate data as described in the previous section for Case 1 and explore all possible combinations of true $\bar{c}_1 = (0.4, 0.7, 0.9)$ and $\theta_0 = \text{logit}(0.01, 0.05, 0.10)$ assuming perfect specificity ($\bar{c}_0 = 0$). We will also explore $\bar{r}$ in $(1, 2, 5, 10)$. This results in a total of 36 simulation settings. Other simulation settings using different values of $\bar{c}_0$, $\bar{c}_1$ and $\bar{r}$ are similar. For each simulation setting, we simulate 100 datasets. For each dataset, we obtain an estimate of $\theta_G^{(simple)}$ and $\theta_0^{(simple)}$. We then use the expressions in **Sections S3.2 and S2.2** to obtain predicted values for $\theta_G$. The true value of $\theta_G$ is 0.5 in all simulation scenarios.

Analysis Intercept is known: We first consider the setting in which $\theta_G^{(simple)}$ and $\theta_0^{(simple)}$ are both available. In this setting, the predictions can be based on (SuppEq. 2.5), which does not vary with $\bar{r}$. In the main paper, we provide boxplots of the predicted values of $\theta_G$ across each of the 100 simulated datasets for potential sensitivity values between 0.1 and 0.9 for a simulation setting in which $\bar{r} = 5$. In each setting, the true sensitivity is denoted with a bolded boxplot, and the true value of $\theta_G$ is illustrated with a horizontal line. The boxplot corresponding to the true value for $\bar{c}_1$ in each simulation setting always contains and is centered somewhat near the true value of $\theta_G$, 0.5. There are some sensitivities for which no solution to (SuppEq. 2.5) exists, as demonstrated in left-most subplots. As mentioned previously, situations where there is no solution correspond to implausible parameter configurations. For some settings (in particular, with small $\theta_0$), there is not much difference in the predicted $\theta_G$ across $\bar{c}_1$. There are two reasons for this. Firstly, low prevalences are associated with low biases in $\theta_G^{(simple)}$ under perfect specificity. Secondly, the true value of $\theta_G$ and resulting $\theta_G^{(simple)}$ are small in these settings, so the absolute bias is expected to be small.

Analysis Intercept is Unknown but Population Prevalence is known: We also consider the setting in which $\theta_0^{(simple)}$ is not known. This may commonly be the case when we use published summary statistics, and many existing pipelines for performing such analyses may not routinely save values for $\theta_0^{(simple)}$. As such, we want to explore plausible values of $\theta_G$ knowing only $\theta_G^{(simple)}$ and a rough value of $\theta_0$ (or a window of values for $\theta_0$). For a given simulated dataset within a particular simulation setting, we can obtain predictions for the value of $\theta_G$ given $\theta_G^{(simple)}$, $\theta_0$ and various potential values of $\bar{c}_1$ and $\bar{r}$ using expression (SuppEq. 2.4). **Figures S3, S4, and S5** provide simulation results for settings in which we have 1%, 5%, and 10% disease rates respectively. In settings where the disease is rare (e.g. less than 1%) we can expect there to be little difference in the predictions across different values of the sensitivity and sampling ratio. Greater differences can be seen when we have larger population disease rates. In all three figures, the median prediction is near the true value of 0.5. This indicates that our expressions have reasonable performance at recovering the true $\theta_G$ through the proposed sensitivity analysis approach.

In practice, we will have a single estimate of $\theta_G^{(simple)}$ rather than 100 values. In **Figure S6**, we demonstrate a sensitivity analysis taking $\theta_G^{(simple)}$ to be the median value of $\theta_G^{(simple)}$ across the 100 simulated datasets. We might expect some additional variability if we use $\theta_G^{(simple)}$ from a single fit. The plotted triangle in this figure indicates the true sampling ratio and $\theta_G$, and the color of the triangle indicates the true sensitivity value. We can see generally good concordance with predicted values of $\theta_G$ and the true value of $\theta_G$ for the true value of $\bar{c}_1$.

#### S7 Additional materials for MGI data analysis

In the main paper, we perform a sensitivity analysis using Michigan Genomics Initiative (MGI) data and exploring phenotype associations with individual genetic loci along with polygenic risk scores. Here, we provide some additional information. Note that we assume $\tilde{c}_0 = 0$ since we believe very few subjects will be incorrectly diagnosed with cancer.

In **(author?)** [4], we compare GWAS results for an EHR-derived breast cancer phenotype in MGI with genotype associations published in the NHGRI-EBI GWAS catalog, which is viewed as a comparative gold standard. We demonstrated that the GWAS results using MGI data are generally similar to results in the GWAS catalog, but there are many specific SNPs for which the results differ, as shown in **Figure S7**.

In the main paper, we explore breast cancer GWAS associations for the 6 particular loci in which (1) the GWAS catalog estimate does not fall into the MGI estimate's confidence interval and (2) the GWAS catalog effect is stronger than the MGI effect in the same direction. We may be interested in performing a similar exploration when the MGI GWAS estimate is very similar to the gold standard. **Figure S8** shows the upper and lower limits of predicted $\theta_G$ for one such SNP. In this example, the MGI estimate and the GWAS catalog estimate are nearly identical.

In the main paper, we also explore PRS-phenotype associations using phenotypes and population prevalences reported in **(author?)** [5] and **(author?)** [4]. **Table S2** reproduces these published values.

#### S8 Obtaining an educated guess for a sampling ratio

The sensitivity analysis explored previously relies on having a rough idea of plausible values for the sampling ratio. However, we often have weak intuition of what this value might be. In this section, we use population disease prevalence estimates to obtain crude values for $r$.

We define "true" $r(Z, \phi) = \frac{\int P(S=1|D=1,W^\dagger,Z)f(W^\dagger|D=1,Z)dW^\dagger}{\int P(S=1|D=0,W^\dagger,Z)f(W^\dagger|D=0,Z)dW^\dagger}$. This value is difficult to pinpoint when $f(W^\dagger|D, Z)$ and $P(S = 1|D, W^\dagger, Z)$ are unknown. However, we may be able to get a rough sense of crude $\tilde{r} = \frac{P(S=1|D=1)}{P(S=1|D=0)}$ as follows:

$$\tilde{r} = \frac{P(S = 1|D = 1)}{P(S = 1|D = 0)} = \frac{P(D = 1|S = 1)}{1 - P(D = 1|S = 1)} \frac{1 - P(D = 1)}{P(D = 1)}$$

Suppose we have a rough sense of the population disease rate, $P(D = 1)$ (seen in **Table S2** for selected cancers in MGI). We also know $P(D^* = 1|S = 1)$ as the prevalence of the disease in our sampled dataset using the potentially misclassified outcome. We can write

$$P(D^* = 1|S = 1) = P(D^* = 1|D = 1, S = 1)P(D = 1|S = 1) + P(D^* = 1|D = 0, S = 1)P(D = 0|S = 1)$$

$$P(D = 1|S = 1) = \frac{P(D^* = 1|S = 1) - P(D^* = 1|S = 1, D = 0)}{P(D^* = 1|S = 1, D = 1) - P(D^* = 1|S = 1, D = 0)}$$

Let $\tilde{c}_1 = P(D^* = 1|D = 1, S = 1)$ be a crude value for the sensitivity $c_1(Z, \beta) = \int P(D^* = 1|D = 1, X^\dagger, Z, S = 1)f(X^\dagger|D = 1, Z)dX^\dagger$ and $\tilde{c}_0 = P(D^* = 1|D = 0, S = 1)$ be a crude value for the false positive rate $c_0(Z, \alpha) = \int P(D^* = 1|D = 0, Y^\dagger, Z, S = 1)f(Y^\dagger|D = 0, Z)dY^\dagger$. Then we can write

$$\tilde{r} = \frac{P(D^* = 1|S = 1) - \tilde{c}_0}{\tilde{c}_1 - P(D^* = 1|S = 1)} \frac{1 - P(D = 1)}{P(D = 1)}$$

We can use this expression to obtain a crude value for $r$ using the population prevalence $P(D = 1)$ and the sample phenotype prevalence $P(D^* = 1|S = 1)$ across different $\tilde{c}_1$ and $\tilde{c}_0$. Under perfect specificity, we can use the above expression, setting $\tilde{c}_0 = 0$.

##### S8.1 Demonstration in MGI and UKB

We consider the disease prevalence rates and MGI sample phenotype prevalences for various cancers as reported in **(author?)** [4] and reproduced in part in **Table S2**. We can use these values to obtain a crude sampling ratio for our dataset. Since we believe the crude sensitivity will be at least 0.5 for these cancers, we restrict our focus to $\tilde{c}_1 >= 0.5$. For this analysis, we assume $\tilde{c}_0 = 0$ since we believe very few subjects will be incorrectly diagnosed with cancer.

The left panel in **Figure S9** shows the results for MGI. Melanoma is predicted to have the largest corresponding sampling ratio among the cancers considered. This is unsurprising as Michigan Medicine is known for its skin cancer treatment center, and many patients come from the surrounding areas for treatment. Overall, the crude sampling ratios in MGI tend to be greater than 1, reflecting the disease enrichment seen in MGI due to the sampling mechanism of recruiting surgical patients within Michigan Medicine.

As a comparison, we perform this same analysis for the UK Biobank, a large population-based biobank with nearly 500,000 participants, many of whom have matched EHR and genetic information. We expect the sampling ratio to be less extreme for UK Biobank than for Michigan Genomics Initiative due to their very different sampling strategies. The right panel in **Figure S9** shows the results for UK Biobank. As expected, the crude sampling ratio values are generally smaller than in MGI. Indeed, the crude sampling ratio values are usually less than 1. This is intuitive– the UK Biobank restricts recruitment to subjects aged 40-69. Since cancer is primarily a disease of older age, many older patients are excluded from entry. Therefore, we might expect UK Biobank to have undersampling of diseased subjects in the population, which we find reflected in our crude estimates for the sampling ratio. These explorations should not be taken as proof of the "true" value for $\bar{r}$. Instead, these explorations are intended to be used as a way to find a reasonable benchmark guiding subsequent sensitivity analyses.

### S9 Allowing for dependence between $G$ and $W$ (avoiding assumption 4)

#### S9.1 Assuming imperfect specificity

Suppose instead we do not make assumption 4, so we allow $W$ to be associated with $G$ given $D$ and $Z$. In this case and assuming imperfect specificity, under **assumptions 1-3**, we have that

$$\text{logit}\left(P(D^*=1|G,Z,S=1)\right)$$
$$=\log\left(\frac{c_0(Z)}{1-c_0(Z)}\right)+\log\left(e^{\theta_0+\theta_G G+\theta_Z Z}\frac{c_1(Z)r(Z,G)}{c_0(Z)}+1\right)-\log\left(e^{\theta_0+\theta_G G+\theta_Z Z}\frac{(1-c_1(Z))r(Z,G)}{1-c_0(Z)}+1\right)$$

where

$$r(Z,G;\phi)=\frac{\int P(S=1|D=1,W^\dagger,Z)f(W^\dagger|D=1,Z,G)dW^\dagger}{\int P(S=1|D=0,W^\dagger,Z)f(W^\dagger|D=0,Z,G)dW^\dagger}$$

We now approximate the above expression using a first order Taylor Series approximation with respect to $Z$ and $G$, where $\bar{Z}$ represents the mean of $Z$ and $\bar{G}$ represents the mean of $G$ in the sample. Let $\bar{c}_1=c_1(\bar{Z})$, $\bar{c}_0=c_0(\bar{Z})$, and $\bar{r}=r(\bar{Z},\bar{G})$. We can write

$$\text{logit}\left(P(D^*=1|G,Z,S=1)\right)\approx\log\left(\frac{\bar{c}_0}{1-\bar{c}_0}\right)+\left\{\frac{1}{\bar{c}_0}+\frac{1}{1-\bar{c}_0}\right\}\left\{\frac{\partial c_0(Z)}{\partial Z}\Big|_{Z=\bar{Z}}\right\}(Z-\bar{Z})$$
$$+\log\left(e^{\theta_0+\theta_G\bar{G}+\theta_Z\bar{Z}}\frac{\bar{c}_1\bar{r}}{\bar{c}_0}+1\right)+\frac{e^{\theta_0+\theta_G\bar{G}+\theta_Z\bar{Z}}\frac{\bar{c}_1\bar{r}}{\bar{c}_0}\theta_G+e^{\theta_0+\theta_G\bar{G}+\theta_Z\bar{Z}}\frac{\bar{c}_1}{\bar{c}_0}\left\{\frac{\partial r(\bar{Z},G)}{\partial G}\right\}\Big|_{G=\bar{G}}}{e^{\theta_0+\theta_G\bar{G}+\theta_Z\bar{Z}}\frac{\bar{c}_1\bar{r}}{\bar{c}_0}+1}(G-\bar{G})$$
$$+\frac{e^{\theta_0+\theta_G\bar{G}+\theta_Z\bar{Z}}\frac{\bar{c}_1\bar{r}}{\bar{c}_0}\theta_Z+e^{\theta_0+\theta_G\bar{G}+\theta_Z\bar{Z}}\left\{\frac{\partial\frac{c_1(Z)r(Z,\bar{G})}{c_0(Z)}}{\partial Z}\Big|_{Z=\bar{Z}}\right\}}{e^{\theta_0+\theta_G\bar{G}+\theta_Z\bar{Z}}\frac{\bar{c}_1\bar{r}}{\bar{c}_0}+1}(Z-\bar{Z})$$
$$-\log\left(e^{\theta_0+\theta_G\bar{G}+\theta_Z\bar{Z}}\frac{(1-\bar{c}_1)\bar{r}}{1-\bar{c}_0}+1\right)-\frac{e^{\theta_0+\theta_G\bar{G}+\theta_Z\bar{Z}}\frac{(1-\bar{c}_1)\bar{r}}{1-\bar{c}_0}\theta_G+e^{\theta_0+\theta_G\bar{G}+\theta_Z\bar{Z}}\frac{1-\bar{c}_1}{1-\bar{c}_0}\left\{\frac{\partial r(\bar{Z},G)}{\partial G}\right\}\Big|_{G=\bar{G}}}{e^{\theta_0+\theta_G\bar{G}+\theta_Z\bar{Z}}\frac{(1-\bar{c}_1)\bar{r}}{1-\bar{c}_0}+1}(G-\bar{G})$$
$$-\frac{e^{\theta_0+\theta_G\bar{G}+\theta_Z\bar{Z}}\frac{(1-\bar{c}_1)\bar{r}}{1-\bar{c}_0}\theta_Z+e^{\theta_0+\theta_G\bar{G}+\theta_Z\bar{Z}}\left\{\frac{\partial\frac{(1-c_1(Z))r(Z,\bar{G})}{1-c_0(Z)}}{\partial Z}\Big|_{Z=\bar{Z}}\right\}}{e^{\theta_0+\theta_G\bar{G}+\theta_Z\bar{Z}}\frac{(1-\bar{c}_1)\bar{r}}{1-\bar{c}_0}+1}(Z-\bar{Z})$$

**Suppose further that the covariate set $Z$ is centered** on the sample such that $\bar{Z}=0$. In this setting, we have that

$$\theta_G^{(simple)}\approx\left[\frac{e^{\theta_0+\theta_G\bar{G}+\theta_Z\bar{Z}}\frac{\bar{c}_1\bar{r}}{\bar{c}_0}}{e^{\theta_0+\theta_G\bar{G}+\theta_Z\bar{Z}}\frac{\bar{c}_1\bar{r}}{\bar{c}_0}+1}-\frac{e^{\theta_0+\theta_G\bar{G}+\theta_Z\bar{Z}}\frac{(1-\bar{c}_1)\bar{r}}{1-\bar{c}_0}}{e^{\theta_0+\theta_G\bar{G}+\theta_Z\bar{Z}}\frac{(1-\bar{c}_1)\bar{r}}{1-\bar{c}_0}+1}\right]\theta_G$$
$$+\left[\frac{e^{\theta_0+\theta_G\bar{G}+\theta_Z\bar{Z}}\frac{\bar{c}_1}{\bar{c}_0}}{e^{\theta_0+\theta_G\bar{G}+\theta_Z\bar{Z}}\frac{\bar{c}_1\bar{r}}{\bar{c}_0}+1}-\frac{e^{\theta_0+\theta_G\bar{G}+\theta_Z\bar{Z}}\frac{(1-\bar{c}_1)}{1-\bar{c}_0}}{e^{\theta_0+\theta_G\bar{G}+\theta_Z\bar{Z}}\frac{(1-\bar{c}_1)\bar{r}}{1-\bar{c}_0}+1}\right]\left\{\frac{\partial r(\bar{Z},G)}{\partial G}\right\}\Big|_{G=\bar{G}}$$

The first term in the above sum is the same as appears in SuppEq. 2.3. The second term expresses the relationships between the sampling ratio and $G$. When $G$ is weakly associated with $W$, the final term $\left\{\frac{\partial r(\bar{Z},G)}{\partial G}\right\}\Big|_{G=\bar{G}}$ may be small, so small violations of **Assumption 4** may not strongly impact results. However, when this derivative is larger, results may be more strongly impacted by the relationship between $G$ and $W$.

We have that

$$\frac{\partial r(\bar{Z}, G; \phi)}{\partial G} = \frac{\partial \left[\frac{\int P(S=1|D=1,W^\dagger,Z)f(W^\dagger|D=1,Z,G)dW^\dagger}{\int P(S=1|D=0,W^\dagger,Z)f(W^\dagger|D=0,Z,G)dW^\dagger}\right]}{\partial G}$$

assuming we can reverse the order of integration and differentiation, we have that

$$\frac{\partial r(\bar{Z}, G; \phi)}{\partial G} = \frac{\left[\int P(S=1|D=1, W^\dagger, Z)\frac{\partial f(W^\dagger|D=1,Z,G)}{\partial G}dW^\dagger\right]}{\left[\int P(S=1|D=0, W^\dagger, Z)f(W^\dagger|D=0, Z, G)dW^\dagger\right]}$$
$$+ r(Z, G)\frac{\left[\int P(S=1|D=0, W^\dagger, Z)\frac{\partial f(W^\dagger|D=0,Z,G)}{\partial G}dW^\dagger\right]}{\left[\int P(S=1|D=0, W^\dagger, Z)f(W^\dagger|D=0, Z, G)dW^\dagger\right]}$$

#### S9.2 Assuming perfect specificity

Suppose instead that we assume $\bar{c}_0 = 0$. We have

$$\begin{aligned}
&\text{logit}\left(P(D^* = 1|G, Z, S = 1)\right)\\
&= \log\left(\frac{P(D=1|G,Z)c_1(Z)r(Z)}{P(D=1|G,Z)(1-c_1(Z))r(Z) + P(D=0|G,Z)}\right)\\
&= \log\left(\frac{e^{\theta_0+\theta_G G+\theta_Z Z}c_1(Z)r(Z)}{e^{\theta_0+\theta_G G+\theta_Z Z}(1-c_1(Z))r(Z)}\right)\\
&= \theta_0 + \theta_G G + \theta_Z Z + \log\left(c_1(Z)\right) + \log\left(r(Z)\right) - \log\left(e^{\theta_0+\theta_G G+\theta_Z Z}(1-c_1(Z))r(Z) + 1\right)
\end{aligned}$$

We now approximate the above expression using a first order Taylor Series approximation with respect to $Z$ and $G$, where $\bar{Z}$ represents the mean of $Z$ and $\bar{G}$ represents the mean of $G$ in the sample. We can write

$$\text{logit}\left(P(D^*=1|G,Z,S=1)\right) \approx \theta_0 + \theta_G G + \theta_Z Z + \log\left(\bar{c}_1\right) + \left[\frac{1}{\bar{c}_1}\right]\left\{\left.\frac{\partial c_1(Z)}{\partial Z}\right|_{Z=\bar{Z}}\right\}(Z-\bar{Z})$$
$$+ \log\left(\bar{r}\right) + \left[\frac{1}{\bar{r}}\right]\left\{\left.\frac{\partial r(Z,\bar{G})}{\partial Z}\right|_{Z=\bar{Z}}\right\}(Z-\bar{Z}) + \left[\frac{1}{\bar{r}}\right]\left\{\left.\frac{\partial r(\bar{Z},G)}{\partial G}\right|_{G=\bar{G}}\right\}(G-\bar{G})$$
$$- \log\left(e^{\theta_0+\theta_G\bar{G}+\theta_Z\bar{Z}}(1-\bar{c}_1)\bar{r} + 1\right) - \frac{e^{\theta_0+\theta_G\bar{G}+\theta_Z\bar{Z}}(1-\bar{c}_1)\bar{r}\theta_G + e^{\theta_0+\theta_G\bar{G}+\theta_Z\bar{Z}}(1-\bar{c}_1)\left.\left\{\frac{\partial r(\bar{Z},G)}{\partial G}\right\}\right|_{G=\bar{G}}}{e^{\theta_0+\theta_G\bar{G}+\theta_Z\bar{Z}}\frac{(1-\bar{c}_1)\bar{r}}{1-\bar{c}_0} + 1}(G-\bar{G})$$
$$- \frac{e^{\theta_0+\theta_G\bar{G}+\theta_Z\bar{Z}}\frac{(1-\bar{c}_1)\bar{r}}{1-\bar{c}_0}\theta_Z + e^{\theta_0+\theta_G\bar{G}+\theta_Z\bar{Z}}\left\{\left.\frac{\partial\frac{(1-c_1(Z))r(Z,\bar{G})}{1-c_0(Z)}}{\partial Z}\right|_{Z=\bar{Z}}\right\}}{e^{\theta_0+\theta_G\bar{G}+\theta_Z\bar{Z}}\frac{(1-\bar{c}_1)\bar{r}}{1-\bar{c}_0} + 1}(Z-\bar{Z})$$

**Suppose further that the covariate set $Z$ is centered** on the sample such that $\bar{Z} = 0$. In this setting, we have that

$$\theta_G^{(simple)} \approx \frac{1}{e^{\theta_0+\theta_G\bar{G}}(1-\bar{c}_1)\bar{r} + 1}\left[\theta_G + \frac{1}{\bar{r}}\left\{\left.\frac{\partial r(\bar{Z},G)}{\partial G}\right|_{G=\bar{G}}\right\}\right]$$

#### S9.3 Allowing $W$ to contain $G$ under perfect specificity

We now consider an extension where sampling directly depends on $G$. This may be the case for general $G$, but may be less common in the setting where $G$ is a SNP or a PRS. In this case, we have the same expression for $\theta_G^{(simple)}$ except this time we have

$$r(Z, G; \phi) = \frac{\int P(S=1|D=1, W^\dagger, Z, G) f(W^\dagger|D=1, Z, G) dW^\dagger}{\int P(S=1|D=0, W^\dagger, Z, G) f(W^\dagger|D=0, Z, G) dW^\dagger}$$

Suppose for simplicity that $W^\dagger$ is empty, so either $G$ and $D$ are the only factors related to sampling or the remaining factors in $W$ are included in $Z$. Then we have

$$r(Z, G; \phi) = \frac{P(S=1|D=1, Z, G)}{P(S=1|D=0, Z, G)}$$

Suppose we model sampling using a logistic regression, where $\text{logit}(P(S=1|D,Z,G)) = \phi_0 + \phi_D D + \phi_Z Z + \phi_G G$. Assuming $\bar{Z} = 0$, we have

$$r(Z, G; \phi) = e^{\phi_D} \frac{1 + e^{\phi_0 + \phi_Z Z + \phi_G G}}{1 + e^{\phi_0 + \phi_D + \phi_Z Z + \phi_G G}}$$

$$\left.\frac{\partial r(\bar{Z}, G)}{\partial G}\right|_{G=\bar{G}} = \frac{\phi_G e^{\phi_0 + \phi_Z \bar{Z} + \phi_G \bar{G}} \left[1 - e^{\phi_D}\right]}{(1 + e^{\phi_0 + \phi_D + \phi_Z \bar{Z} + \phi_G \bar{G}})^2} = \frac{\phi_G e^{\phi_0 + \phi_G \bar{G}} \left[1 - e^{\phi_D}\right]}{(1 + e^{\phi_0 + \phi_D + \phi_G \bar{G}})^2}$$

and

$$\theta_G^{(simple)} \approx \frac{1}{e^{\theta_0 + \theta_G \bar{G}}(1 - \bar{c}_1)\bar{r} + 1} \left[\theta_G + \frac{e^{\phi_0 + \phi_G \bar{G}}}{1 + e^{\phi_0 + \phi_G \bar{G}}} \frac{\phi_G \left[e^{-\phi_D} - 1\right]}{1 + e^{\phi_0 + \phi_D + \phi_G \bar{G}}}\right]$$

We note that the second term of the above expression is exactly zero if $\phi_D = 0$. However, if $\phi_D \neq 0$, $\phi_0$, $\phi_D$, and $\phi_G$ all contribute to the bias.

#### S10 Bias for rare diseases

Suppose we are interested in a disease that is very rare in the target population, e.g. a disease rate of 1/2000 patients. We are interested in understanding the expected *relative* biases in these settings.

Suppose first that we have perfect specificity. In this case, we have that

$$\theta_G^{(simple)} \approx \left[\frac{1}{e^{\theta_0+\theta_G\bar{G}}(1-\bar{c}_1)\bar{r}+1}\right]\theta_G$$

Unless $\theta_G$ is very large, we have that $e^{\theta_0+\theta_G\bar{G}} \approx 0$. We have that $\theta_G^{(simple)} \to \theta_G$ as $\theta_0 \to -\infty$. Therefore, we expect very little relative bias in $\theta_G^{(simple)}$ in the perfect-specificity rare disease setting unless $\theta_G$ is very large for fixed values of $\bar{c}_1$ and $\bar{r}$. That being said, we could still have large absolute bias even with small relative bias if $\theta_G$ is very large. This may be the case for some rare, highly penetrant SNPs.

Suppose, however, that we have imperfect specificity strictly less than 1. We also assume that sensitivity is strictly less than 1, which we expect to be the case for EHR data. In this case, we have

$$\theta_G^{(simple)} \approx \left[\frac{\frac{\bar{c}_1}{\bar{c}_0}-\frac{(1-\bar{c}_1)}{1-\bar{c}_0}}{\left[e^{\theta_0+\theta_G\bar{G}}\frac{\bar{c}_1\bar{r}}{\bar{c}_0}+1\right]\left[e^{\theta_0+\theta_G\bar{G}}\frac{(1-\bar{c}_1)\bar{r}}{1-\bar{c}_0}+1\right]}\right]\theta_G e^{\theta_0+\theta_G\bar{G}}\bar{r}$$

and $\theta_G^{(simple)} \to 0$ as $\theta_0 \to -\infty$ for fixed values of $\bar{r}$, $\bar{c}_0$, and $\bar{c}_1$. This implies that our simple analysis will tend to produce null results when we have imperfect sensitivity and specificity for rare diseases given $\bar{r}$, $\bar{c}_0$, and $\bar{c}_1$.

We note that with rare diseases, we might expect $\bar{r}$ to be large, particularly if we are using case-control sampling based on that rare disease. Suppose instead we consider very large values of $\bar{r}$. Suppose we have $\theta_0 \approx \text{logit}(1/2000) \approx -7.6$. **Figure S10** shows the degree of shrinkage toward the null for different combinations of $\bar{r}$, $\bar{c}_0$, and $\bar{c}_1$. This plot demonstrates that, even for a very small fixed $\theta_0$ and imperfect specificity, we may not expect perfect shrinkage to the null if the sampling ratio is very large. In contrast, under perfect specificity (where we might expect little relative bias for rare diseases) we see increased relative (and absolute) bias for very large sampling ratios. This strange phenomenon is a function of the disease rate in the *sample*. If we have a very large sampling ratio, then even for a rare disease the disease may be very well-represented in the sample. Imperfect specificity will, therefore, have a weaker impact on the amount of outcome misclassification than if the sampling ratio were 1.

**Table S1:** *Bias in target model parameter $\theta_G$ in the simple analysis allowing for imperfect specificity*

| Variable | Value | **Specificity = 1** ($\bar{c}_0 = 0$) | **Specificity < 1** ($\bar{c}_0 > 0$) |
|---|---|---|---|
| Sensitivity ($\bar{c}_1$) | Decreases toward 0 | Bias increases | Bias increases |
| | • No Underreporting ($\bar{c}_1 = 1$) | No Bias | Biased |
| | • Underreporting ($\bar{c}_1 < 1$) | Biased | Biased |
| Sampling Ratio ($\bar{r}$) | Increases toward $\infty$ | Bias increases if $\bar{c}_1 < 1$ | Bias may increase or decrease |
| | • Sampling Independent of $D$ ($\bar{r} = 1$) | Biased if $\bar{c}_1 < 1$ | Biased |
| | • Sampling Dependent on D ($\bar{r} \neq 1$)$^\dagger$ | Biased if $\bar{c}_1 < 1$ | Biased |
| Disease Prevalence | Increases toward 1 | Bias increases if $\bar{c}_1 < 1$ | Bias may increase or decrease |
| Absolute Effect of $G$ | Increases toward $\infty$ | Bias increases if $\bar{c}_1 < 1$ | Bias may increase or decrease |

$^\dagger$ We consider sampling dependent on observed case-control status, $D^*$, in **Section S4**

**Table S2:** *PRS-phenotype associations along with MGI and US prevalences for selected cancers from* **(author?)** *[4] and* **(author?)** *[5], based on 30,702 unrelated patients of recent European ancestry in MGI*

| Cancer Type | PRS OR (95% CI) | MGI Prevalence | Lifetime Disease Rate in US |
|---|---|---|---|
| Colorectal | 1.3 (1.1, 1.6) | 2.6 | 4.2 |
| Breast (female) | 2.3 (2.0, 2.7) | 12.4 | 12.4 |
| Melanoma of skin | 2.4 (2.0, 2.8) | 6.2 | 2.3 |
| Prostate (male) | 3.3 (2.7, 3.9) | 12.4 | 11.2 |
| Bladder | 1.4 (1.2, 1.7) | 3.7 | 2.3 |
| Non-Hodgkins Lymphoma | 1.3 (1.1, 1.6) | 3.1 | 2.1 |

**Figure S1:** *Structure of data with two sampling steps*

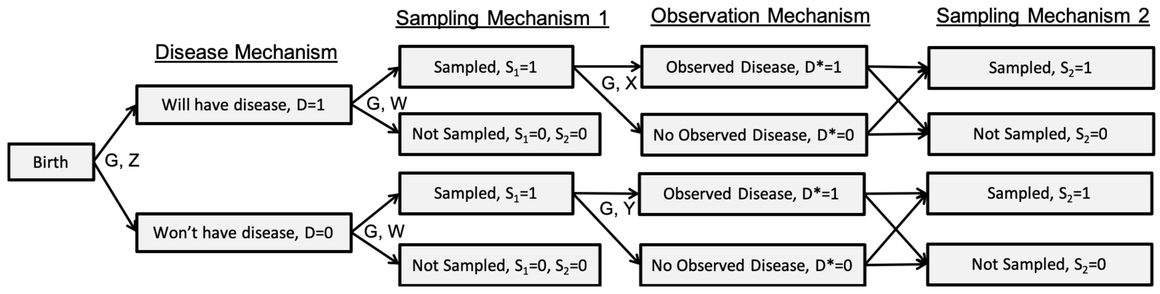

**Figure S2:** *Correspondence between estimated and approximated values of* $\theta^{(simple)}$. Each point corresponds to a simulation setting. Simulation estimates have been averaged across 50 ($G$ is a PRS) or 100 ($G$ is a SNP) simulated datasets.

(a) $\theta_0^{(simple)}$ when $G$ is a SNP

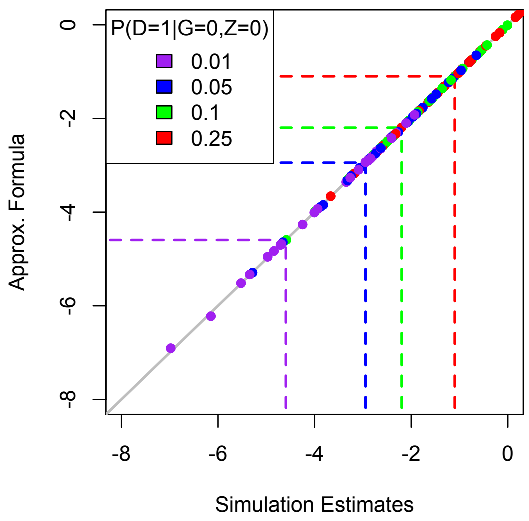

(b) $\theta_G^{(simple)}$ when $G$ is a SNP

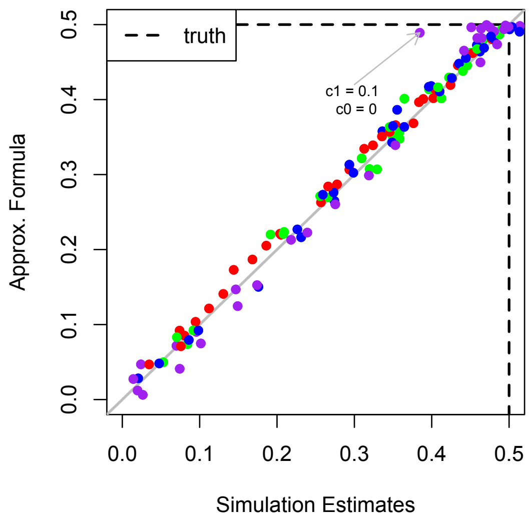

(c) $\theta_0^{(simple)}$ when $G$ is a PRS

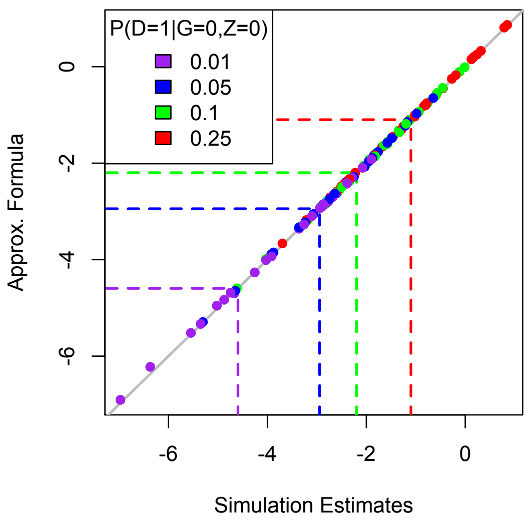

(d) $\theta_G^{(simple)}$ when $G$ is a PRS

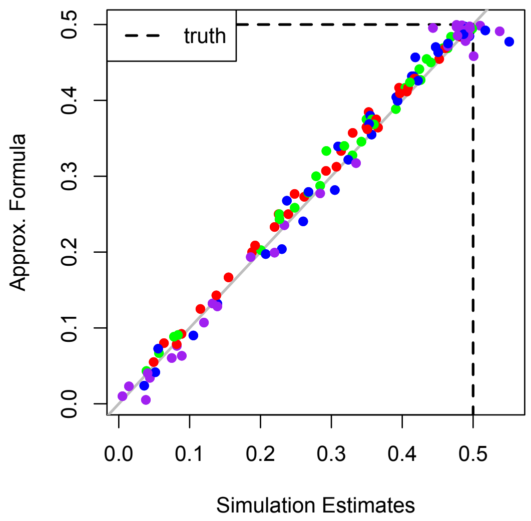

**Figure S3:** *Predicting $\theta_G$ for different sensitivities and sampling ratios assuming known 1% disease rate in population* Inter-quartile range for predicted $\theta_G$ across simulated datasets. The black horizontal line indicates the true value of $\theta_G = 0.5$. The bolded IQR indicates the true sensitivity.

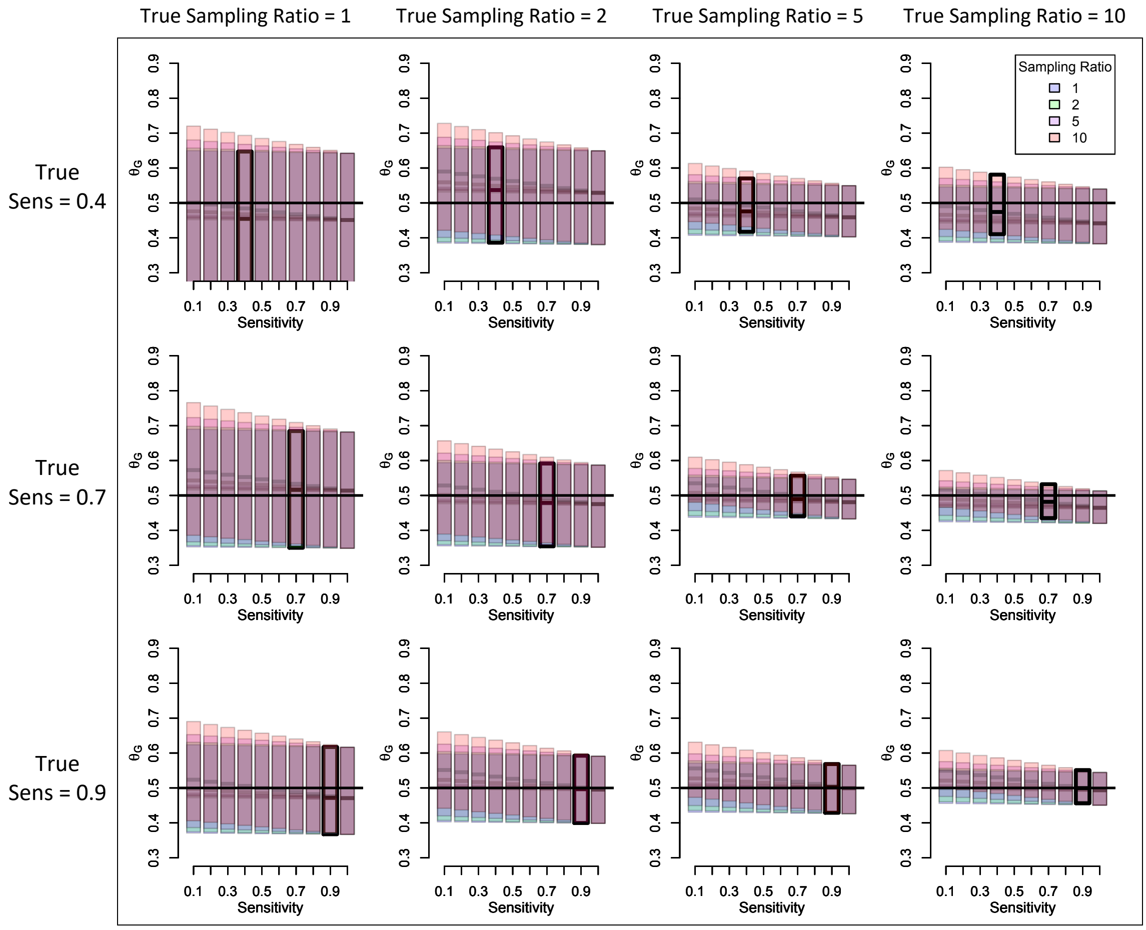

**Figure S4:** *Predicting $\theta_G$ for different sensitivities and sampling ratios assuming known 5% disease rate in population* Inter-quartile range for predicted $\theta_G$ across simulated datasets. The black horizontal line indicates the true value of $\theta_G = 0.5$. The bolded IQR indicates the true sensitivity.

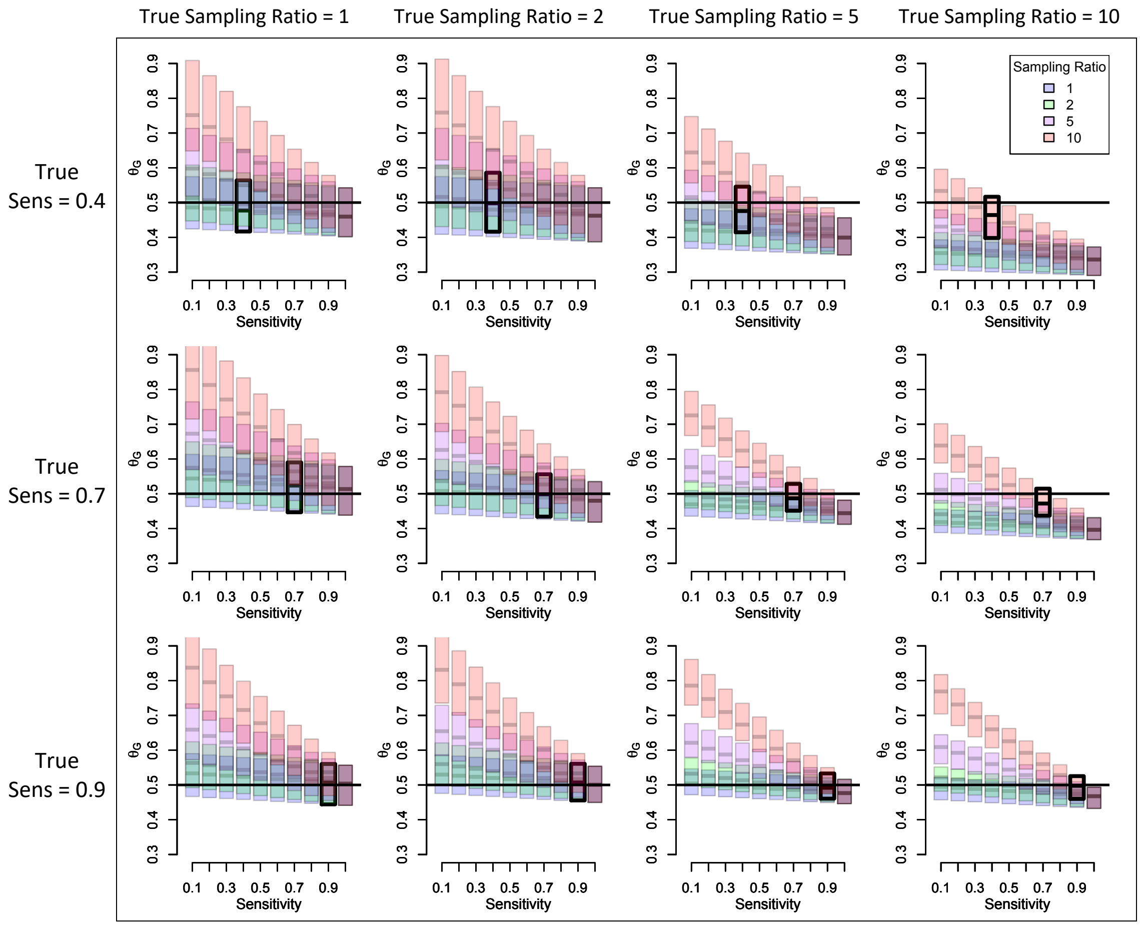

**Figure S5:** *Predicting $\theta_G$ for different sensitivities and sampling ratios assuming known 10% disease rate in population* Inter-quartile range for predicted $\theta_G$ across simulated datasets. The black horizontal line indicates the true value of $\theta_G = 0.5$. The bolded IQR indicates the true sensitivity.

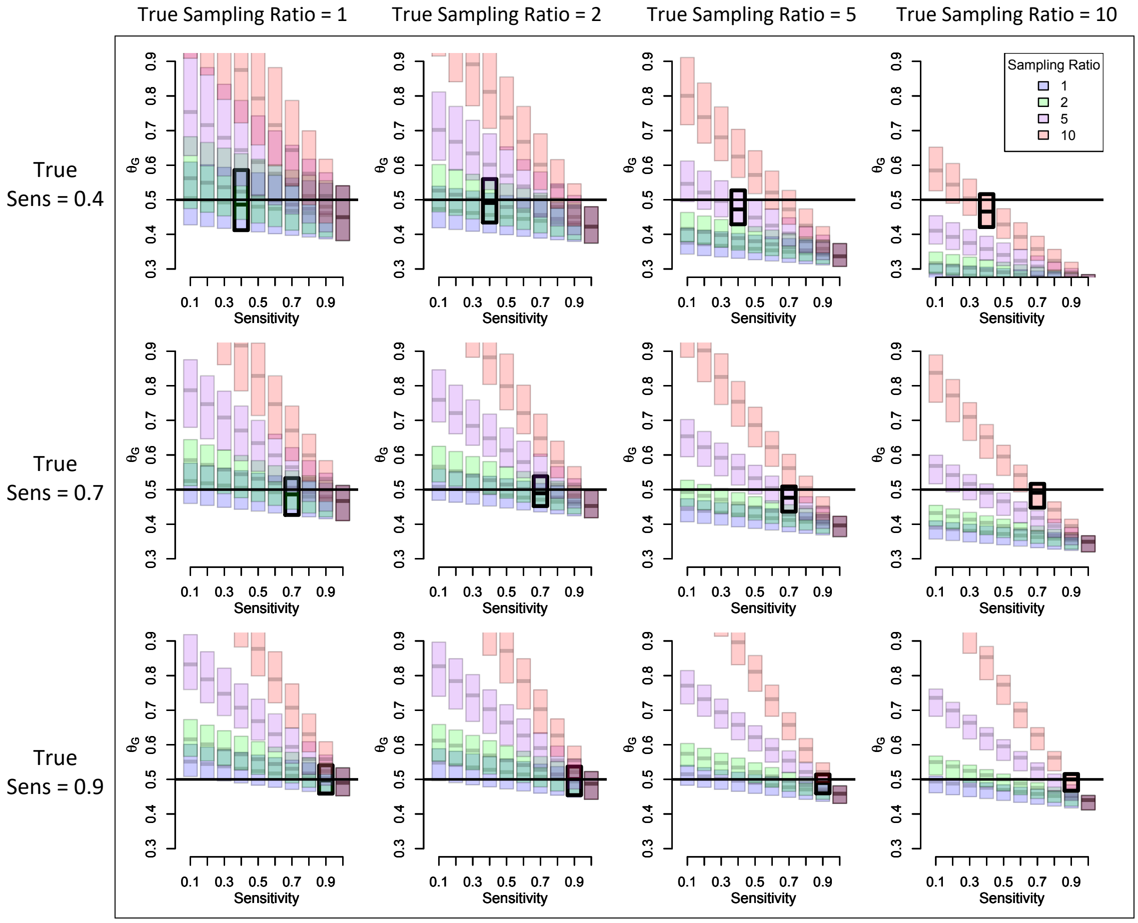

**Figure S7:** *NHGRI-EBI GWAS catalog breast cancer GWAS results ($\theta_G$) compared to GWAS results based on over 38,000 unrelated patients of recent European ancestry in MGI ($\theta_G^{(simple)}$).* Confidence intervals for MGI GWAS estimates are shown in gray.

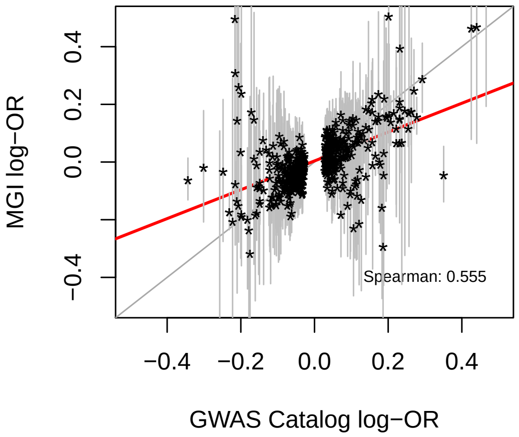

**Figure S6:** *Predicting $\theta_G$ for different sensitivities and sampling ratios using a single value for $\theta_G^{(simple)}$* The color and location of the triangle corresponds to true values for the sensitivity and sampling ratio.

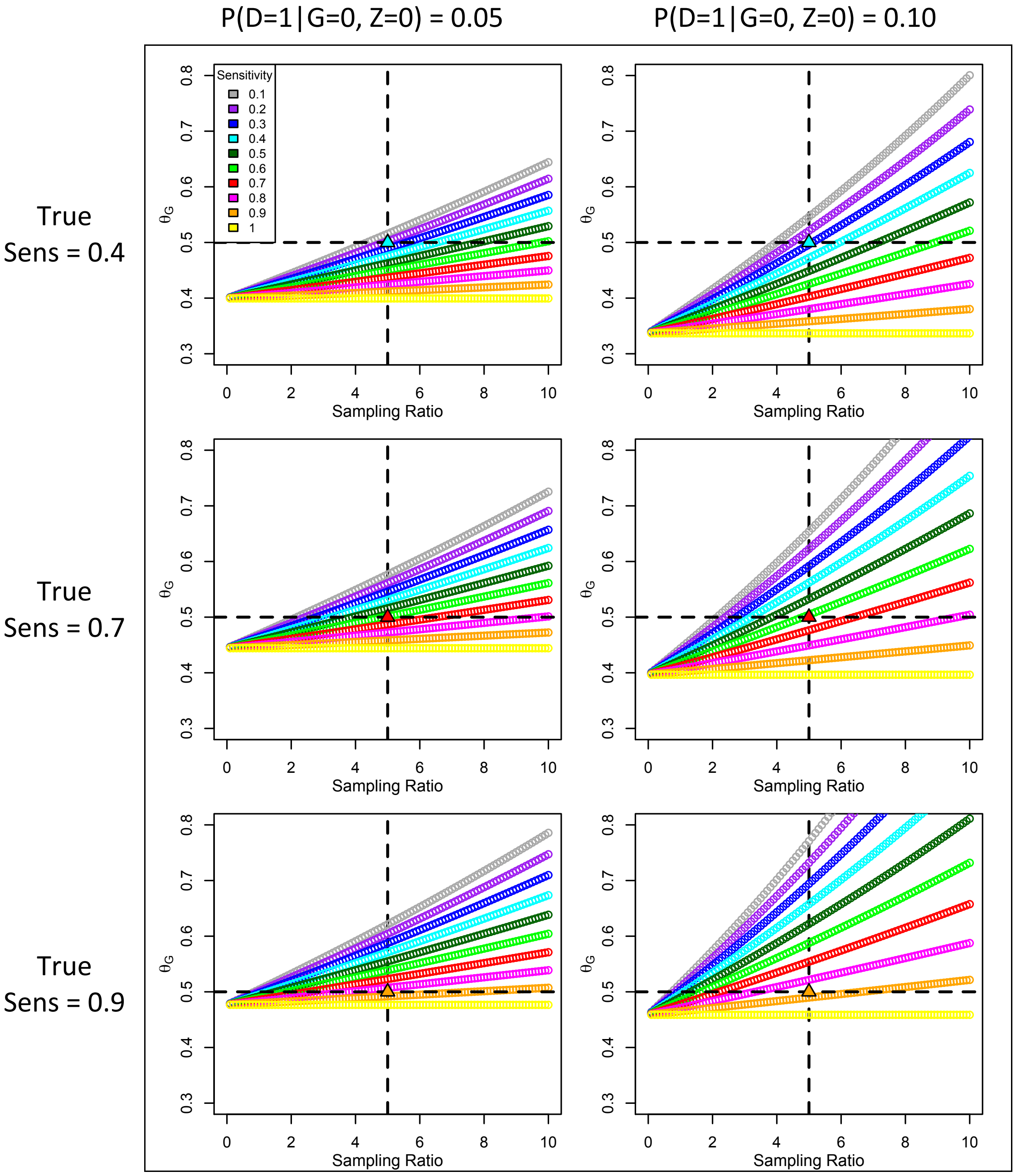

**Figure S8:** *Plausible values for $\theta_G$ in MGI for a breast cancer association in which MGI and the GWAS catalog agree.* The NHGRI-EBI GWAS catalog (Michigan Genomics Initiative) estimate and corresponding 95% confidence interval is shown in black (gray).

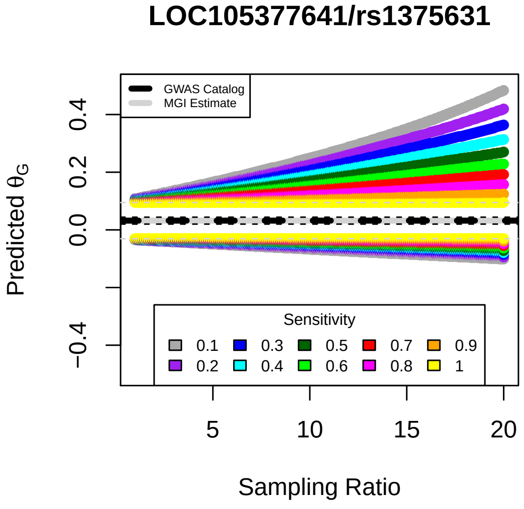

**Figure S9:** *Obtaining plausible values for the sampling ratio.* The horizontal dotted line corresponds to a sampling ratio of 1

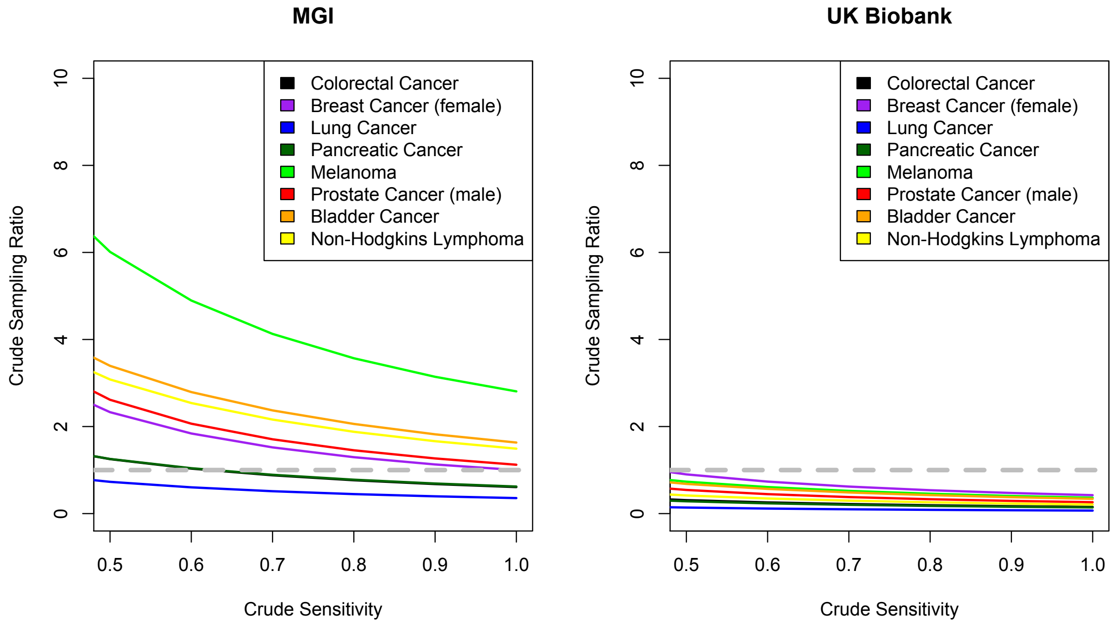

**Figure S10:** Degree of relative bias for rare disease with prevalence = 1/2000

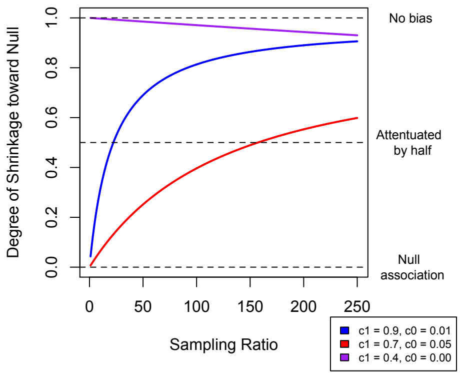
